# Exploration of endo-xylanase from novel strain of *Bacillus velezensis* AG20 isolated from the cave of Meghalaya

**DOI:** 10.1101/2020.04.06.028878

**Authors:** Arabinda Ghosh, Shravanika Mahanta, Subhro Banerjee, Debabrat Baishya

## Abstract

Cave sets the example of extreme ecological niche and habitat for diversified microorganisms. Present study involved in the isolation of endoxylanase producing novel strain *Bacillus velezensis* AG20 from the Krem Phyllut cave, Meghalaya, India. Culture dependent studies, molecular phylogentics, RNA secondary folding pattern based on 16S rDNA substantiated the identity of this novel strain. *Bacillus velezensis* AG20 revealed the superbug quality having resistance against various class of broad-spectrum antibiotics. *Bacillus velezensis* AG20 revealed biofilm formation over the cell surface in FESEM. Highest cell biomass and xylanase production supported in TB medium, further purified partially to 5.3 fold with 21% yield. Molecular weight of the purified xylanase found to be 45 kDa. Enzyme kinetics and pattern of hydrolysis revealed the evidence for the selection of linear birchwood xylan with V_max_ = 21.0 ± 3.0 U/ml, Km = 1.25 mg/ml, K_cat_ = 1.75/s at optimum pH 7 and temperature 50°C also found significant statistically in Taguchi’s orthogonal design. Conversely, ruled out any exo-acting activity against synthetic pNP-xylopyranoside substrate. Endo-xylanase isolated from *Bacillus velezensis* AG20 was moderately thermostable over temperatures 50 and 60°C. Time dependent hydrolysis of agro-waste sugar cane bagasse depicted the production of xylooligosaccharides (XOS) predominantly xylobiose, xylotriose and xylotetrose. Purified mixed XOS hold their prebiotic potential by promoting the growth of probiotics *Bifidobacterium* and *Lactobacillus* as well as high stability (~90%) against systemic fluids. Mixed XOS (300 μg/ml) displayed anti-proliferation activities by reducing the growth of HT-29 and Caco-2 cells significantly 90% and 75%, respectively, after 48 h.

**IMPORTANCE:** Extremophiles dwelling inside the caves have laden with the extraordinary capabilities of bioconversion by nature. The pristine ecological niche inside the cave, absence of proper light and air, supports the livelihood of novel microorganisms. In India, Meghalaya is hoisting longest caves in the East Khasi Hills, providing conducive environment for novel bacterial strains. With the prime objective of isolating novel bacterial strains that produce extracellular xylanase our studies have been carried out. Considering the present industrial demand for nutraceutical, prebiotics, anti-proliferating agents and biofuels by the conversion of lignocellulosic biomass (LCB), novel enzymes are required. Xylanases from bacterial origin play a significant role in conversion of LCB into oligosaccharides. Therefore, exploration and characterization of xylanase producing novel isolate from cave may pave the new arena for the production of prebiotic and anti-inflammatory oligosaccharides from agro-waste.

## Introduction

Cave is a pristine and ecologically extreme environment for life. In the interior of caves, the light source is scanty and therefore the phototrophic microorganisms are unable to synthesize organic matter. World’s highest precipitation, combination of quality limestone, low elevation, and a hot and moist climate have occasioned in the formation of Meghalaya’s subterranean caves. The three hills, Khasi, Jaintia, and Garo, contain limestone of variable quantity and quality (1). Favourable conditions inside the cave facilitate to host diverse microorganisms in the dim natural light and artificially lighted subterranean environments that group themselves into biofilms associated with rock surfaces (2). These biofilms are complex aggregates of microorganisms embedded in a self-produced matrix that provides protection for growth, enabling microorganisms to survive adverse cave environments (2). The primary productivity of caves relies upon the chemolitho-autotrophic microorganisms (3). The majority of the cave dwelling microorganisms are oligotrophic in nature (4). The Mawsmai cave and Krem Phyllut caves in East Khasi hills, Meghalaya, India is the habitat for many important microorganism in the speleotherms (5). Their investigation reported to have microfabric features of speleotherms showing strong resemblance with fossilised bacteria. Molecular techniques further prevailed the confirmation of *Bacillus cereus, Bacillus mycoides, Bacillus licheniformis, Micrococcus luteus,* and Actinomycetes in the speleotherms (5). From the caves Krem Soitan, Krem Mawpun, and Krem Lawbah, 113 bacterial isolates were identified and majority belonged to the genus *Bacillus, Rummeliibacillus, Staphylococcus,* and *Brevibacterium* (6). In another report Banerjee and coworkers reported the abundance of the genera *Bacilllus* and *Pseudomonas* from Krem Mawsmai, Krem Mawmluh and Krem Mawjymbuin caves of East Khasi Hills, Meghalaya, India (7). Not many reports have been entailed since then about the geo-microbial aspects from the caves of Meghalaya, India. *Bacillus* strains are well pronounced for their biotransformation capacity in the caves through a number of processes, including photosynthesis, ammonification, denitrification, sulphate reduction, enzymatic hydrolysis and anaerobic sulphide oxidation (6, 7, 8). *Bacillus* sp. is a potential source of xylanases among bacteria, and a number of bacilli, such as *B. stearothermophilus, B. circulans, B. licheniformis, B. pumilus, B. subtilis, B. halodurans* and *B. amyloliquefaciens* attributed to synthesize extracellular xylanase (9, 10, 11).

Xylan is the most abundant non-cellulosic polysaccharide, found to be of approximately 20–35%, thus as a whole comprises approximately one third of all renewable organic carbon sources on earth. In annual plants and hardwoods, xylan is the most abundant non-cellulosic polysaccharide which accounts for 20-35% of the total dry weight in biomass (12, 13). Depending upon the plant origin, the xylan backbone becomes covalently bind to varying degree of acetic acid, arabinose, D-glucuronic acid, 4-O-methyl-D-glucuronic acid, ferulic acid and p-cumaric acid (14). Apart from the three basic chemical compounds (cellulose, hemicellulose, and lignin), lignocellulosic biomass contains water, proteins, minerals and other compounds. The organic component of biomass plays a major role in processing and producing biofuel (15). Chemical hydrolysis of lignocelluloses results in hazardous by-products, prospecting the use of microbial enzymes, which are species in action for xylan hydrolysis and is an environment friendly option. Xylanase (endo-1, 4-β-xylanase) and β-xylosidase (β-D-xyloside xylohydrolase) are the major constituents of well known microbial xylanolytic enzyme systems. Cellulases and xylanases have an assortment of industrial applications including pulp & paper, laundry, food, animal feed, brewery and wine, textile, bioenergy industry etc. Endo-β(1→4)-xylanase (E.C 3.2.1.8) belongs to family 3, 5, 8, 10, 11, 30, 43, 51, 98 and 141 glycoside hydrolases (GHs) categorized on the basis of protein folds and hydrolysis of xylan substrates (http://www.cazy.org/). In order to make the enzyme applications more cost effective at industrial level, its production using low cost substrates such as agro-wastes, food and industrial waste, fruit waste, vegetable waste and weed plants as a low-cost solid substrate for the production of xylanases. Xylanases have potential to overcome the problem of high cost enzyme production and recommended by many workers (16, 17, 18). The costs of enzyme production can be reduced by optimizing the fermentation process such as medium, temperature and pH, which are the basic goals of research for industrial applications (19).

In our earlier study *Bacillus licheniformis* DM5 was explored from a hotspring and investigated for the production of extracellular xylanase. Nevertheless, that investigation complied the production of xylo-oligosaccharides (XOS) from agro-waste corncob (11). Till date xylanase belongs to family GH10 and GH11 predominantly involve in xylo-oligosaccharide production (Javier et al., 2018). Arabino-xylooligosaccharides (AXOS) and XOS gained much interest owing to their prebiotic potential and raw material from agro industrial waste. In these processes, extracellular xylanases were used from *Bacillus halodurans* S7, *Bacillus subtilis, Clostridium thermocellum* ATCC 27405, *Geobacillus thermodenitrificans* TSAA1. Enzymes belong to GH10 protein family were utilized to hydrolyze wheat straw, wheat bran sugar cane bagasse, wheat spelt xylan, brich wood, beech wood xylan (21, 22, 23, 24). XOS act as the prebiotic, which can be used as ingredients of functional food, cosmetics, pharmaceuticals or agricultural products and as a plant growth regulator. In addition to the health effects, XOS present interesting physico-chemical properties, they are moderately sweet, and stable over a wide range of pH and temperatures and have organoleptic characteristics suitable for incorporation into foods (25, 26, 27). XOS have importance in decreasing the blood lipids, protecting liver functions, decreasing blood pressure, anticancer and regulating blood sugar. Therefore, XOS-containing diets are considered beneficial in improving gastrointestinal health.

Our present study involves the isolation of novel endo-xylanse bacterial strain from a virgin cave Krem Phyllut situated in the East Khasi Hills of Meghalaya, India. Of late, no such strong evidences on isolation and characterization of extracellular xylanase producing bacteria from the cave samples are reported which may be beneficial for fulfilling the need of industry in future. Moreover, we have also investigated for their potential application in XOS production from agro industrial waste and to exploring their potentials to become prebiotic and anti-prolific activities.

## MATERIALS AND METHODS

### CHEMICALS AND REAGENTS

Nutrient Agar, nutrient broth, luria bertani broth, muller hinton agar, glucose, dextrose, sucrose, lactose, mannitol, phenol red, starch agar, Gram’s iodine, yeast extract, gelatin, skim milk powder, agar, peptone, tryptone, kovac’s reagent, mr-vp media, simmons citrate agar, nutrient agar, nutrient broth, safranin, crystal violet, vp reagent (i), vp reagent (ii), oxidase disc, ferrous ammonium sulphate, sodium thiosulphate, potassium dihydrogen phosphate, Bradford’s reagent, DNS reagent, hydrogen peroxide, petroleum ether, glacial acetic acid, n-Butanol were procured from Himedia Laboratories Pvt. Ltd. India. Trypsin, bile salts, Taq polymerase, dNTPs, α-Amylase procured from Sigma Aldrich, USA.

### SAMPLE COLLECTION

Soil sample was collected from the virgin cave Krem Phyllut, located in Sohra (Cherrapunjee), East Khasi Hills, Meghalaya: N 26°09.202’ & E° 91°39.633’, elevation: 1169 metres were measured by using a GPS system (Garmin ETrex Touch 35, Netherlands). Ambient temperature inside the cave was measured to be 18°C, soil pH 6.8 and the relative humidity 93%. Cave floor sediment samples have collected in sample container and maintained at 18°C for further analyses. The experimental analyses have been further carried out in the Microbial Ecology Laboratory, Department of Botany and Bioenergy and Bioprospective Laboratory, Department of Bioengineering and Technology, GUIST, Gauhati University.

### ISOLATION OF BACTERIAL CULTURES

One gram of soil sediment sample was diluted into 90 ml distilled water and dilutions (10^-3^, 10^-4^, 10^-5^, 10^-6^, 10^-7^), were prepared separately. Then 0.1 ml of suspension from each dilution has transferred into nutrient agar and R2A agar plates by using spread plate and pour plate technique and incubated at 37°C for 24 h. Morphologically distinct colonies selected and pure cultures were prepared by repeated streaking on nutrient agar plates and slants and incubated at 37 °C for 24 h.

### MORPHOLOGICAL AND BIOCHEMICAL TESTS

The bacterial isolates were then characterized morphologically, physiologically and biochemically following standard protocol as described by Cappuchino and Sherman (28). Bacterial isolates were subjected to gram staining to analyze morphologically and physiologically. Biochemical tests viz. i) Fermentation of sugars: glucose, mannitol, lactose and sucrose;, ii) hydrogen sulphide production test, iii) citrate utilization test, iv) Sugar utilization test was performed with Hi-Carbohydrate Kit^TM^ (Himedia, India), v) catalase and vi) oxidase tests were performed to detect the preliminary characteristics of the isolated bacterial colonies.

### ANTIBIOTIC SUSCEPTIBILITY TEST

Antibiotic susceptibility test was performed by disc diffusion assay following the method of Kirby Bauyer (29). In this study commercial antibiotic discs *viz.* Amoxicillin (10 μg), Ampicillin (10 μg), Cefalexin/cephalexin (30 μg), Cephalothin (30 μg), Chloramphenicol (30 μg), Clindamycin (2 μg), Cloxacillin (5 μg), Co-trimoxazole (25 μg), Erythromycin (15 μg), Gentamicin (10μg), Oxacillin (1 μg), Penicillin-G (2 Units), Tetracycline (10 μg), Vancomycin (10 μg) and Kanamycin (5 μg) were used against the isolated bacterial cultures. Muller Hinton Agar plates were prepared as per the manufacturer’s instruction. The isolates were inoculated on the agar plates and spreaded uniformly. Antibiotic disc placed on the agar plates and incubated at 37 °C for 48 h. A control was kept without antibiotics. The zone of inhibition has been measured using a Vernier calliper.

### SCREENING OF BACTERIAL ISOLATES FOR EXTRACELLULAR XYLANASE ACTIVITY

In-vitro screening of bacterial isolates producing extracellular xylanase was performed by congo red plate assay method as described elsewhere (30). The bacterial isolates were cultured in 1% (w/v) beech wood xylan containing LB agar plates at 37 °C, overnight. After the bacterial enumeration into unilayer, the culture plates were flooded with 0.1% (w/v) congo red solution and left for 15 min with intermittent shaking. Following the congo red staining, plates were washed with distilled water to remove any additional unbound stain and finally washed with 1M NaCl solution to distain the plates. Counter staining wasperformed with 1N HCl for better understanding of the xylanase producing zones.

### MOLECULAR AND PHYLOGENETIC ANALSIS OF EXTRACELLULAR XYLANASE PRODUCING ISOLATE

Molecular phylogenetics facilitates the reconstruction of evolutionary relationships between two closest neighbours for identification of organisms by using macromolecular sequences. To establish the evolutionary relationship and molecular identification of bacteria, genomic DNA (gDNA) from cultivated xylanase producing colony was extracted using gDNA isolation kit (HiPurA™ Bacterial Genomic DNA Purification Kit, Himedia Pvt. Ltd, India). Bacterial cells were harvested after culturing in LB media at 37 °C, 180 rpm overnight and 1.5 ml of 12 h culture was spun down and re-suspended in recommended buffer and the extraction procedure was followed as described in the kit. The conserved 16S rDNA sequence was amplified in PCR reaction using 1492R-5’-TACGGYTACCTTGTTACGACTT-3’ and 27F-5’-AGAGTTTGATCMTGGCTC AG-3’ universal primers and followed by Sanger sequencing at Eurofin Pvt. Ltd., Bangalore, India. The sequenced products were used for the identification of bacterial isolates as well as to find the evolutionary relationship between the samples. In order to make a single contig of the sequencing results DNA baser tool was used. The contig was subjected to BLASTn (http://www.ncbi.nlm.nih.gov/BLASTn) with default parameters for 16S rDNA sequence amongst the bacterial clan to find the sequence homology with the representative members. Sequences with 100–99% query coverage, E-value 0.0 and homology above 95% were considered as closest homologue in the BLASTn output result for multiple sequence alignment and the phylogenetic relationship were established by constructing a phylogenetic tree using MEGA X v 10.1.7.

After the molecular phylogeny analysis, the species specificity of the bacterial isolate was identified further by predicting the 16S rRNA secondary fold structure and the minimum free energy (MFE) among the closely related organisms (11). Secondary folding structures were predicted and validated using MFOLDv3.6 (31) and Vienna RNA web service (http://rna.tbi.univie.ac.at/), respectively, at 37 °C and salt concentration 1M (NaCl) and no divalent cation (Mg^2+^ = 0 M). Kinetic and thermodynamic parameters were calculated using Oligo-Calc open source (http://biotools.nubic.northwestern.edu/ Oligo Calc.html).

### FESEM ANALYSIS OF THE BACTERIAL ISOLATE

Broth culture 1.5 ml was centrifuged at 13,000 rpm for 10-12 minutes. The supernatant discarded and the pellet was resuspended in MB grades of alcohol for dehydration (v/v, 10%, 30%, 50%, 70%, 90%, 100%). The pellet was then picked and a smear was prepared on a coverslip, subsequently dried and coated with gold film using a SC7620 ‘‘Mini’’, Polaron Sputter Coater (Quorum Technologies, Newhaven, England). Field emission scanning electron microscopy (FESEM) image of bacterial colony producing high extracellular xylanase was analyse in electron microscope (FESEM-Carl Zeiss, SIGMA VP-300, Austria).

### CONDITION OPTIMIZATION, PRODUCTION AND PURIFICATION OF XYLANASE

Selected colonies showing exo-xylanase activity in plate assay were cultured in different media containing 1% (w/v) beech wood xylan in order to obtain higher production of xylanase. Four different sets of media were prepared viz. Luria Bertani (LB), 5xLB, Auto inducing Medium (AIM) and Terrific Broth (TB medium) were prepared as described elasewhere (11, 32). Each 100 ml culture broth inoculated with the bacterial colony showing exo-xylanse activity and incubated at 37 °C, 180 rpm overnight. Cells were then harvested from 100 ml culture broth by centrifugation at 13,000 rpm for 10 min and the supernatant separated. Cell free supernatant containing proteins was subjected to 80% (w/v) ammonium sulfate precipitation till the saturation achieved at 4 °C. The partially purified protein was dialyzed against 50 mM sodium phosphate buffer at 4 °C overnight. Partially purified protein was observed in 12% SDS PAGE. The concentration of the partially purified protein in each medium from 100 ml culture was estimated using Bradford’s reagent.

### ENZYME ACTIVITY ASSAY

Initially, the enzyme assays of partially purified xylanse were carried out in 100 μl reaction mixture in 50 mM sodium phosphate buffer pH 7.0 containing 1% (w/v) beech wood xylan. 10 μl of enzyme incubated with 1% (w/v) birchwood xylan at 50°C for 15 min. Temperature optimization assays were carried out between 20°C and 100°C, respectively, for 15 min. Optimization of pH was carried out in 50 mM sodium phosphate buffer between 5.8 to 7.8 following the similar methods as described earlier (11). The enzyme activity was measured by estimating the liberated reducing sugar by dinitrosalicylic acid (DNS) method as described elsewhere. The mixture was diluted by adding water to make up the volume to 1 ml and the absorbance at 540 nm (A_540_) was measured by a UV-Visible. D-Xylose in the range of 10-500 μg/ml was used for generating the standard plot. One unit (U) of enzyme activity is defined as the amount of enzyme that liberates one μmole of reducing sugar per min. The assays of xylanase with synthetic substrates p-nitrophenyl glycoside (pNP-glycosides) viz., p-nitrophenyl-β-D-xylopyranoside and p-nitrophenyl-α-D-xylopyranoside were carried out by estimating the release of 4-nitrophenol (pNP) at 405 nm using a UV-Vis spectrophotometer (Agilent, G 68608) as described elsewhere (11).

The enzyme activity based on physiological conditions has statistically optimized using Taguchi’s orthogonal array. Three independent variables temperature, pH and buffer ionic concentrations were investigated to disseminate the significance of enzyme activity in the evaluation Minitab v 19. Taguchi’s method has been widely accepted in optimization analysis and is a powerful design. The taguchi method operates systematically with fewer trials, thus reducing the time, cost and effort and offers more quantitative information. In this application, L-27 (3^3^) orthogonal matrix was used to study the relationship between the physicochemical factors on the purified xylanase activity. Each of the above selected conditions were taken at three defined levels covering the range over which its effect can be determined. With the objective to find the optimization parameters to maximize the enzyme activity. For individual run, the S/N ratio conforming to larger-the-better primarily function. Therefore, signal-to-noise ratio function designed by Taguchi design was used given as follows:

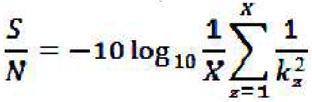

Where, ‘kz’ is signal and ‘X’ is repetitions number during each experiment.

S/N ratios were analysed to find out the main effects of the factors independently; thereafter determining the factors that were statistically significant, by analysis of variance technique (ANOVA). To ascertain the ratio (F) and the p value (p< 0.05), the parameters (factors) in the experimental design at 95% confidence limit were accounted to be statistically significant.

### SUBSTRATE SPECIFICITY OF XYLANASE AGAINST XYLOSE POLYSACCHARIDES

The enzyme activity of xylanase was determined by using 50 mM sodium phosphate buffer at optimum pH and temperature incubated for 15 min. The 100 μl reaction mixture contained 1 % (w/v) substrate, 50 μl of partially purified xylanase. The assays were performed in triplicate. The concentration of reducing sugar was estimated using a standard curve of D-Xylose as partially purified protein has xylanase activity. 100 μl of reaction mixture was taken for estimation of enzyme activity which was calculated against beech wood xylan, birch wood xylan, arabinoxylan, glucuronoxylan. After analysis of substrate specificity the enzyme assays, kinetic parameters viz. K_m_, V_max_ and K_cat_ were determined.

### THERMAL STABILITY STUDY OF THE PURIFIED XYLANASE

Thermal stability is an important parameter for industrially important enzymes for their utilization at elevated temperatures. In this study the temperature optimization of the partially purified xylanase was assessed at varying temperatures from 50°C to 100°C. The partially purified protein (50mM sodium phosphate buffer, pH 7.0) was incubated individually prior to addition of substrate at 50 °C, 60 °C, 70 °C, 80 °C, 90 °C, and 100°C, for 30 min in a water bath. The best activity of the enzyme was tested by taking the OD of the samples at 540 nm.

### EFFECT OF METAL IONS AND DETERGENT ON ENZYME ACTIVITY

Effects of different metal cations, chaotropic agents, and detergent on the activity of partially purified enzyme were determined. The enzyme activity was determined in the presence of various metal salts, such as Ni^2+^ (NiSO_4_· 6H_2_O), Zn2+ (ZnSO_4_·7H_2_O), Cu^2+^ (CuSO_4_·5H_2_O), Co^2+^ (CoCl_2_· 6H_2_O), Mn^2+^ (MnCl_2_·4H_2_O), Al^3+^ (AlCl_3_·6H_2_O), Ca^2+^ (CaCl_2_· 2H_2_O), Fe^3+^ (FeCl_3_.2H_2_O), chaotropic agents like disodium EDTA, urea, guanidine hydrochloride and detergent such as SDS. The assays of partially purified enzyme were performed at 50°C, using 50 mM sodium phosphate buffer (pH 7.0). One-hundred microliters of the reaction mixture containing carob galactomannan (1%, w/v) and metal salt at concentrations (up to 80 mM) or SDS (up to 20 mM) were incubated for 10 min, and a control sample in the absence of the additive was also run. The assays were conducted in triplicates. Partially purified enzyme were incubated with EDTA and urea for 1 h, before measuring the residual activity. The enzyme activity was determined, as described earlier.

### PREPARATION AND PRETREATMENT OF LIGNOCELLULOSIC BIOMASS (LCB)

The sugarcane bagasse was collected, dried at 60°C for 24 h. The fully dried biomass is then grinded and a dry powdery substrate was obtained. The substrate was then pre-treated by Alkaline Peroxide (H_2_O_2_). It is an effective method for pre-treatment of Lignocellulosic biomass (LCB). The powdered LCB was soaked with H_2_O_2_ at room temperature, then the alkaline treatment provided with 40% NaOH (pH of 11-12) at 60°C for a period of 2 h as reported earlier by Kim (33). This method was envisaged for significant removal of lignin (about 70-90%) reported earlier by Singh and Satapathy (15) and the total recovery of xylan content was 64% (w/w). The delignified biomass wass then filtered and allowed to dry for 24 h. The powder obtained then sieved through 0.5 mm mesh and used as substrate for further analysis. Pre-treated and untreated biomass was analysed in FESEM for better understanding of the surface morphology.

### PRODUCTION OF XYLO-OLIGOSACCHARIDE FROM SUGAR CANE BAGGASSE

The qualitative analysis of hydrolyzed products was performed by thin-layer chromatography (TLC) on silica gel-coated aluminium foil (TLC Silica gel 60 F254 20×20 cm, Merck) for detecting sugars. The enzyme with 1% (w/v) birchwood xylan and the LCB was prepared, 100 μl reaction mixtures were incubated at optimized temperature of 60°C and optimized pH 6.9, for 24 h. The reaction products were then dissolved in alcohol and separated from the solid substrate mixture. Then the samples were dried at 60 °C and 0.2 μl of samples were loaded on the TLC plate and kept in the developing chamber saturated with the developing solution (mobile phase) which consisted of acetic acid-n-propanol-water-acetonitrile (4:10:11:14). At the end of the run, migrated sugars were visualized by immersing the TLC plate in a visualizing solution (sulphuric acid/methanol 5:95, v/v; α-napthol 5.0 %, w/v). The TLC plates were then dried at 80°C for 20 min. The migrated spots appeared on the TLC plate. Each TLC spot was then extracted from the silica gel in methanol by centrifugation at 10,000 g for 10 min. Spotted products were extracted in preparative TLC and analyzed using a mass spectrophotometer (Agilent 6550 iFunnel Q-TOF LC/MS system, Agilent 1200 series, USA). The fractions were diluted to 5 ppm in methanol and filtered through a 0.45 μm membrane. ESI–MS in positive ionization mode were carried out for specific fractions and analyzed by TOF detector (Agilent 1200 series, USA).

### APPLICATION OF MIXED OLIGOSACCHARIDES AS PREBIOTICS AND ANTI-PROLIFIC AGENT

To become a potential prebiotic agent, oligosaccharides must bear some significant characteristics such as 1) promotion of growth of human gut probiotic microbiota, 2) should be able to withstand gastric juice, intestinal fluid and oral enzymes (34). In order to perpetuate as growth promoter, in vitro prebiotic potential of the mixed oligosaccharides were investigated on the growth of probiotc strains. Human gut probiotic type strains *Bifidobacterium infantis* NRRL B-41661, *Bifidobacterium longum* NCC 2705 and *Lactobacillus acidophilus* NRRL B-4495 as well as non-probiotic enteric bacteria *E. coli* DH5α and *E. aerogenes* MTCC 3030 were investigated in presence of commercial prebiotic inulin and purified mixed XOS. The purified XOS were pooled and a mixture was prepared for the investigation. Probiotic and human gut non probiotic gut microbes were cultured using enriched and basal medium as described elsewhere (34, 35). Here each growth medium for specific culture of microorganism supplemented with 1% (w/v) inulin, considered as positive control and mixed XOS as test prebiotics. Each culture medium (100 ml) were seeded with 1×10^7^ cells/ml *Bifidobacterium infantis* NRRL B-41661, 3×10^8^ cells/ml *Bifidobacterium longum* NCC 2705, 1×10^8^ cells/ml *Lactobacillus acidophilus* NRRL B-4495, 2×10^6^ cells/ml *E. coli* DH5α and 2×10^8^ cells/ml *Enterobacter aerogenes* MTCC 3030 and were grown at 37°C for 18 h.

Effect of gastric juice on the mixed XOS was examined and compared with the commercial prebiotic inulin for reference. The simulated gastric juice has been prepared (pH 1-4) following the method described elsewhere (34, 36). Freeze-dried XOS mixture was diluted in MlliQ water (pH 5.3) to prepare 1% (w/v) test sample and similarly the reference inulin. The separate reaction mixtures of XOS and inulin with gastric juice were kept at 37°C, 6 h. Samples were picked up in every 1 h interval till 6 h of incubation and the reducing sugar as well as total carbohydrate content estimated using phenol sulphuric acid method as described elsewhere (37, 38). Extent of hydrolysis (%) was estimated using the following equation (36):

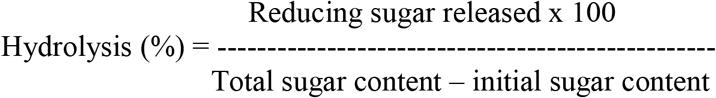

The artificial human intestinal fluid was prepared and its effects were estimated on mixed XOS and reference inulin. Artificial human intestinal fluid was prepared (1,000 U/mL of trypsin and 0.5 % bile salt at pH 8) following the method as described elsewhere (Fernandez et al., 2003). Mixed XOS and inulin were diluted in MlliQ water (pH 5.3) to prepare 1% (w/v) test sample and similarly the reference inulin. The separate reaction mixtures of XOS and inulin with gastric juice were kept at 37 °C, 6 h. Samples were picked up in every 1 h interval till 6 h of incubation and percentage of hydrolysis was estimated similarly as mentioned in the previous method. Effects of α-amylase was estimated on mixed XOS and inulin as described earlier (39). Similar procedures were followed for sample collection and estimation of percentage of hydrolysis as described in the previous method.

In vitro anti-prolific activity of mixed XOS was studied on the human colon cancer cell lines HT-29 and Caco-2. Effect of mixed XOS was examined in a colorimetric assay using 3- (4,5-dimethylthiazolyl-2)-2,5-diphenyltetrazolium bromide (MTT) assay as described elsewhere (40). Colon cancer HT29 and Caco-2 cells were seeded in 96-well plates at density approximately 1.5×10^4^ cells/well and 1.9×10^4^ cells/well incubated at 37°C for 16 h in 5% CO2 atmosphere for surface attachment. Followed by incubation, the media was removed and the mixed XOS at different concentrations were added. A concentration range (1–300 μg/mL) of mixed XOS was investigated. Media without XOS as used as negative control. The plates were again incubated at 37 °C in 5% CO2 atmosphere for 48 h. After 48 h the media were removed and 100 μL MTT (500 μg/mL) was added to each well and further incubated at 37°C for further 4 h. The supernatant was removed and 100 μL dimethylsulfoxide (DMSO) was added to each well. The absorbance at 570 nm was recorded in a 96-well microplate reader (Tecan, Infinite 200 Pro, Tecan Trading AG, Switzerland).

## RESULTS

### MORPHOLOGICAL, BIOCHEMICAL AND ANTIBIOTIC SUSCEPTIBILITY TESTS

The resultant bacterial load of the soil sample collected from the Krem Phyllut Cave of Meghalaya was observed to be 1.8×10^-7^CFU/ml. A total of nine bacterial colonies were selected based on their distinct plate morphological characters and were then isolated in pure form and were designated as PL1, PL2, PL3 PL4, PL5, PL6, PL7, PL8, PL9. The biochemical analyses were performed for the preliminary identification of the isolates. Out of the nine isolates PL1, PL4 and PL7 displayed gram positive, endospore forming and rest were gram negative. From the biochemical tests it could be presumed that all the gram positive and endospore forming isolates may belong to the genus *Bacillus*. Out of the three gram positive isolates only PL1 was observed to utilize xylose during the sugar fermentation test. Moreover, PL1 was efficient in utilizing maltose, dextrose, trehalose, salicin, arabinose, citrate and malonate. Antibiotic susceptibility tests on the nine isolated colonies are depicted in Table 1. PL1 displayed resistance to all the commercial antibiotics used in this experiment. While PL4 showed mixed response of resistance, intermittent and sensitive, PL7 was found to be mostly sensitive against the antibiotics used (Table 1).

**Table 1:**
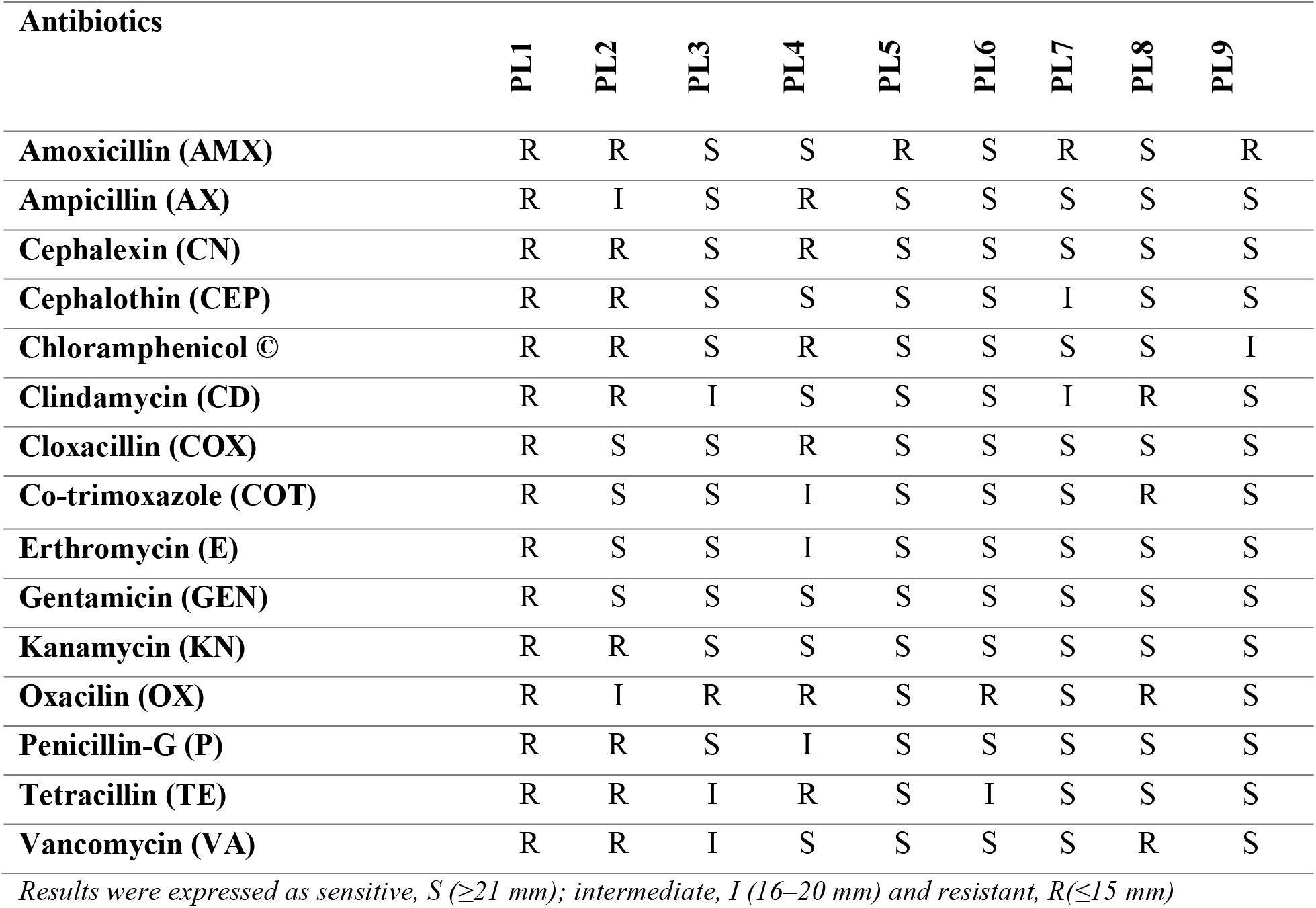
Antibiotic susceptibility tests (disc diffusion assay) on the isolated colonies.

### SCREENING OF BACTERIAL ISOLATES FOR EXTRACELLULAR XYLANASE ACTIVITY

Colonies PL1, PL4 and PL7 in the plate assay with 1% (w/v) beech wood xylan, displayed positive xylanolytic activity by producing a white halo around the colonies. PL1 displayed highest halo zone of xylanolytic activity. This result suggested that this isolate has the capacity to produce extracellular xylanase. Therefore, PL1 was further analysed for the molecular phylogenetics analysis and extracellular xylanase production.

### MOLECULAR CHARACTERIZATION AND PHYLOGENETIC ANALYSIS OF EXTRACELLULAR XYLANASE PRODUCING ISOLATE

Molecular phylogeny anlasysis of the xylanase producing strain PL1 is depicted in figure 1A. 16S rDNA of PL1 displayed 98.9% similarity query coverage 98% and E value 0.0, with the novel bacterium *Bacillus velezensis* Strain NIBSM-PR1 in BLASTn local alignment. Molecular phylogeny based on evolutionary history to find the relatedness among the representative members showed that the PL1 belongs to the species *Bacillus velezensis* on the same branch of the phylogenetic tree (Figure 1A). Although PL1 was found to be closely assosciated with repesentative members *viz. Bacillus* sp, *B. amyloliquefaciens*, divergence times for all branching points in the topology were calculated using the Maximum Likelihood method and Tamura-Nei model in the time tree displayed sharp relatedness with *B. velezensis* (Figure 1B). The marginal difference in similarity, based on mophological and phylogenmetic analsysis the PL1 was designated as *Bacillus velezensis* AG20. Physical and kientic parameters of the colosely associated representative members with *Bacillus velezensis* AG20 diplayed in table 2. 16S rRNA secondary folding pattern analysis exhibited that PL1 has similar arrangements of loops and pseudoloops with *Bacillus velezensis* with minimum free energy (MFE) of −205 and −207 Kcal/mol, respectively (Figure 2A and 2B), which corroborates the MFE of the consensus −209 Kcal/mol (Figure 2C). Conversely, conformational divergence as well as sharp MFE (−160 Kcal/mol) differences were observed also in case of *Bacillus amyloliquefaciens* (Figure 2D). Therefore, it can be suggested that the xylanase producing PL1 was a new bacterial strain *Bacillus velezensis* AG20. Interestingly, it was observed from the table 2, *Bacillus velezensis* AG20 has high GC content (52%) and melting temperature strongly correlated (p < 0.05) the adaptability at the extreme environment in side the cave.

**Figure 1.**
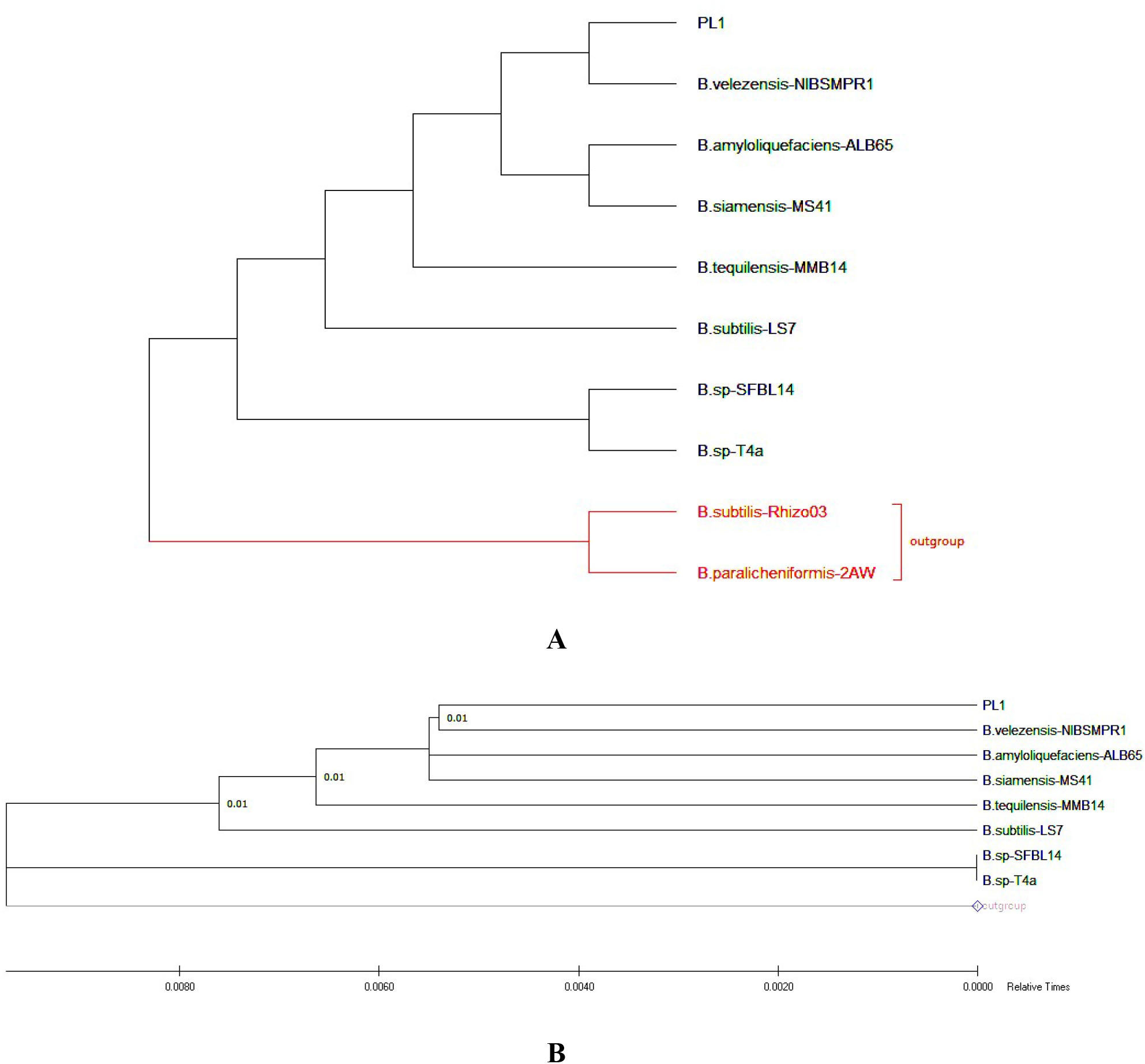
(A) Phylogenetic tree (Maximum likelihood) showing the comparative study of PL1 with 16S rDNA sequence of nine representative members *Bacillus* sp. The 16S rDNA of the isolate from the cave Krem Phyllut displayed closest homology with seven representative strains *Bacillus velezensis* NIBSMRP1 followed by *Bacillus amyloliquefaciens* ALB65, *Bacillus siamensis* MS41, *Bacillus tequilensis* MMB14, *Bacillus subtilis* LS7, *Bacillus* sp. SFBL14, *Bacillus* sp. T4a and two outgroup *B. subtilis* Rhizo03 and *B. paralicheniformis* 2AW (Red). (B) Time tree constructed based on molecular clock for PL1 with the representative members of *Bacillus* to showing the evolutionary pattern and relatedness. The scale bar corresponds to the relative time.

**Figure 2.**
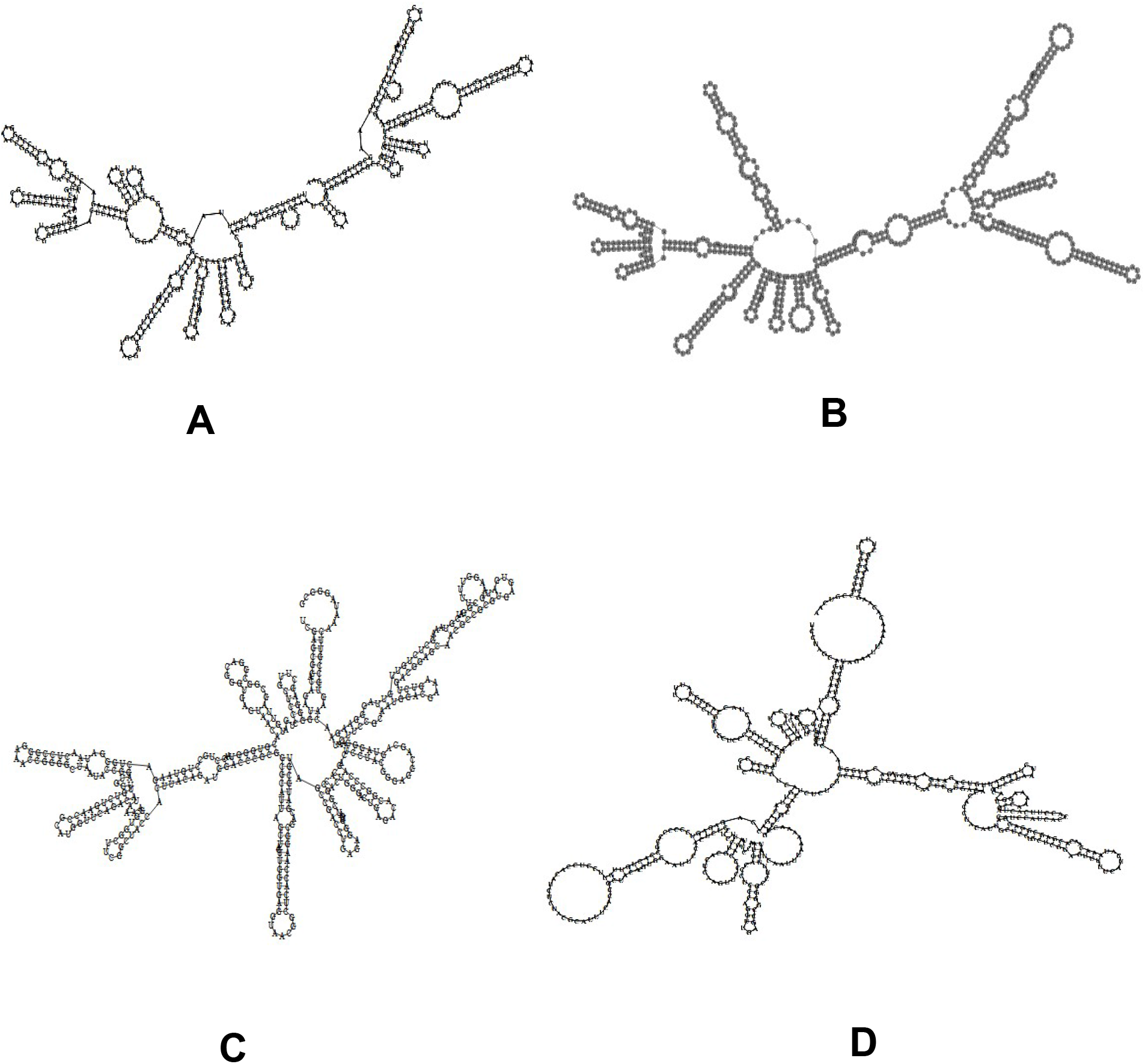
MFOLD 3.6 predicted RNA secondary structures of (A) PL1, (B) *Bacillus velezensis* NIBSMRP1, (D) *Bacillus amyloliquefaciens* ALB65, (C) RNAalifold predicted consensus of PL1 and *Bacillus velezensis* NIBSMRP1.

**Table 2.**
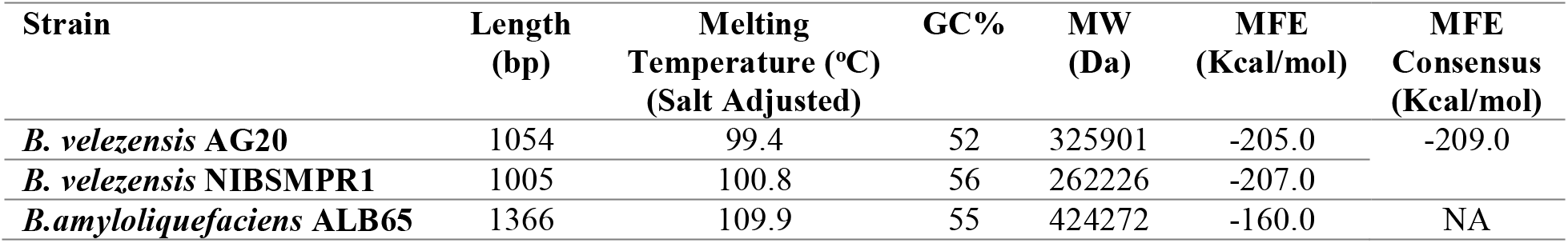
Physical and kinetic parameters of closely related 16S RNA of representative strains with *B. velezensis* AG20

### FESEM ANALYSIS OF *Bacillus valezensis* AG20

Electron micrograph of 14 h culture of *Bacillus velezensis* AG20 was captured and illustrated in figure 3A and 3B. At 1μm scale, 10.00 KX magnification the bacterial cells were appeared to be well separated and distinct entity (Figure 3A). In a much closer view at 50.00 KX magnification and 100 nm scale the single cell of xylanase producing *Bacillus velezensis* AG20 exhibited a smooth thin film covered over cell the surface (Figure 3B). The thin film is due to the production of biofilms which is an usual phenomenon of bacteria and might be due better adaptibility and sustenance under the pristine ecological distribution inside the cave.

**Figure 3.**
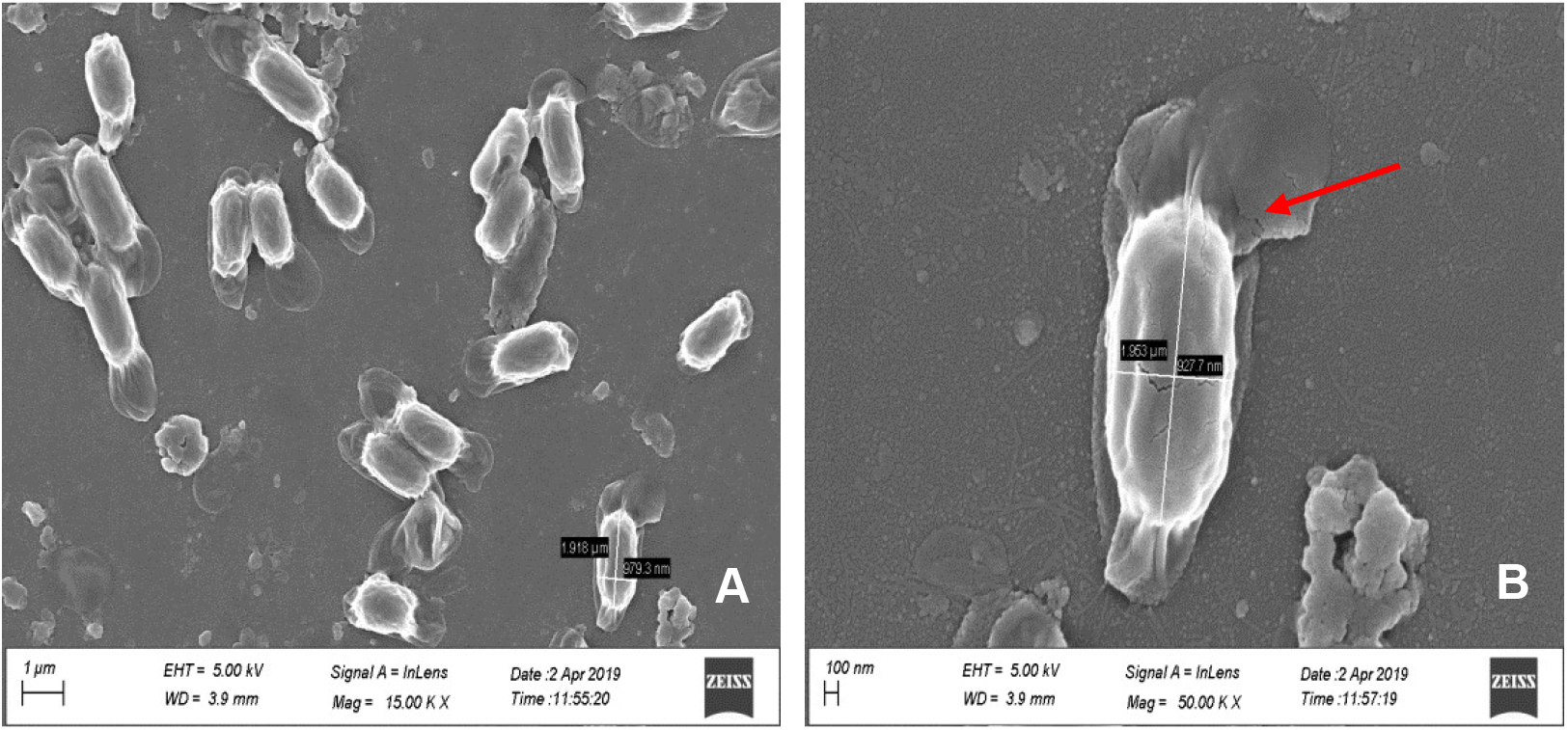
FESEM analysis of *Bacillus velezensis* AG20 (A) bacterial cells displaying in 1 μM scale 15.00 KX resolution, (B) single cell displaying a sheet of membranous biofilm (arrow) over the surface.

### CONDITION OPTIMIZATION, PRODUCTION AND PURIFICATION OF XYLANASE

In order to obtain higher production of extracellular xylanase from *B. velezensis* AG20 media screening was performed. TY medium achieved highest dry cell weight (DCW) 12 g/l and the total protein content using Bradford’s protein estimation method was obtained to be 400 mg/l. Whereas, in AIM the cell biomass was obtained to be 8 g/l and the total protein concentration 302 mg/l. In contrast, low concentrations of cell biomass and protein concentrations were obtained in LB and 5XLB medium 3g/l, 100 mg/l and 1 g/l, 75 mg/l, respectively. The DCW and the protein concentrations in the other related media were measured as the following order TB > AIM >LB > 5xLB (Table 3). Therefore, TB medium was considered for further enzyme production experiments. The total protein was partially purified from the TB medium and the fold purification was obtained to be 5.3 with 21% yield. The partially purified protein displayed a molecular weight of 45 kDa (Figure 4A) in 12% SDS-PAGE. Partially purified xylanase activity was determined using 1% (w/v) commercial substrates viz. birchwood, beechwood, Glucuronoxylan and arabino xylan. The partially purified xylanase showed maximum activity 21.0 ± 3.0 U/ml against birchwood xylan, 19 ± 2.0 U/ml with beechwood xylan, 11.0 ± 1.0 U/ml with Glucuronoxylan and 4.12 ± 0.4 U/ml with arabino xylan. Optimization of temperature and pH were also determined for the purified xylanase. Elevation of incubation temperature from 20 °C to 100 °C, the enzyme activity of xylanase with birchwood xylan displayed an elevation profile till 50 °C where the maximum activity observed 19.91 ± 4.0 U/ml. The activity started falling after 50 °C and at 100 °C just 2% activity was retained (Figure 4B). Therefore, the optimum temperature for the enzyme activity was found to be 50°C. pH optimization experiment revealed that the partially purified xylanase showed maximum activity at pH 7.0, followed by a fall thereafter (Figure 4C). Enzyme activity was found at this pH was 20.8 ± 2.0 U/ml. Nevertheless, the activity was well retained till pH 7.8 (~20%). Partially purified extracellular xylanase did not show any activity against pNP-β-D-xylopyranoside and with pNP-α-D-xylopyranoside. Based on the enzyme activity against natural as well as synthetic substrates, it was evident that the partially purified enzyme is predominantly an endoxylanase.

**Figure 4.**
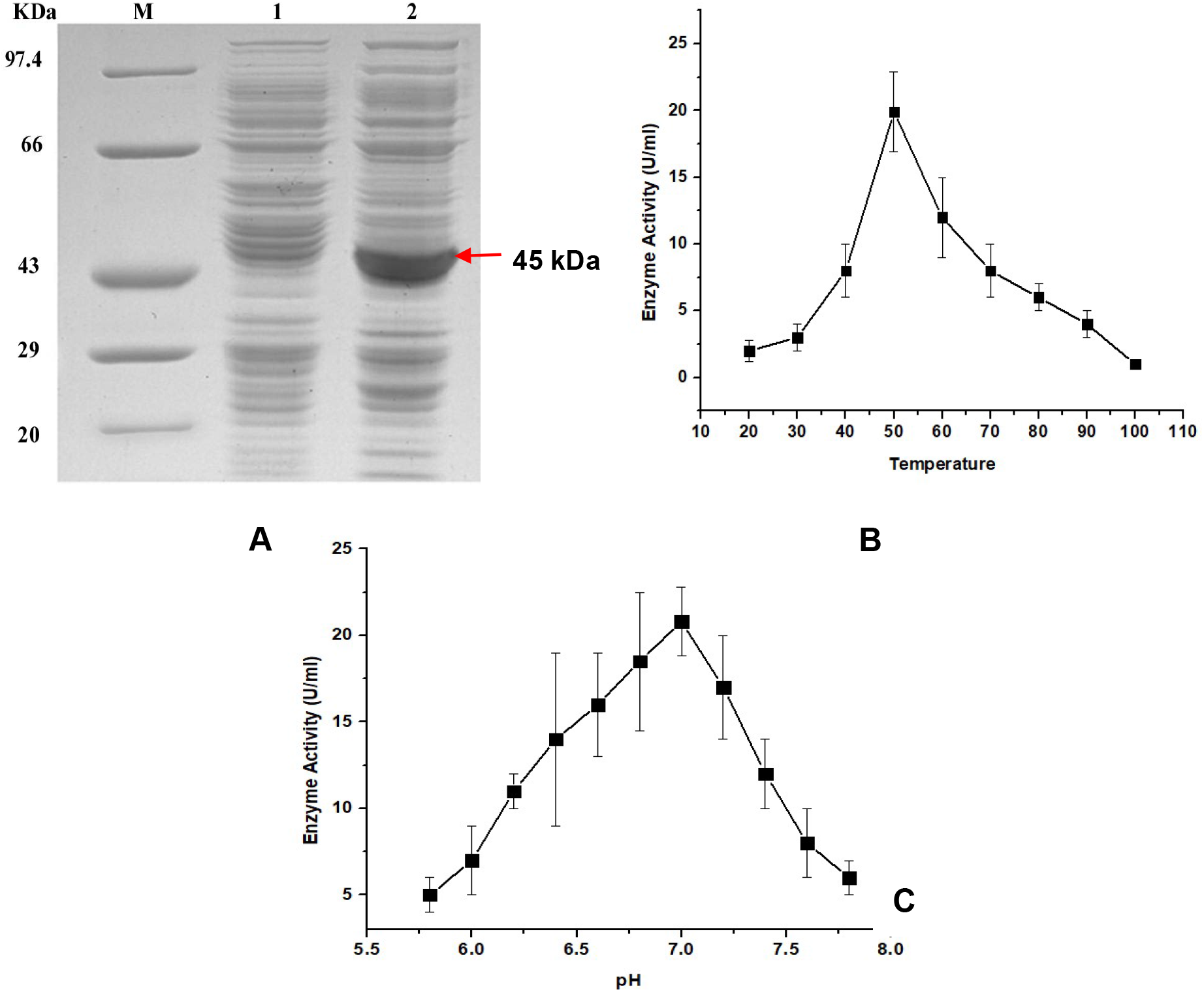
(A) 12% SDS-PAGE of partially purified xylanase from *Bacillus velezensis* AG20 [Lane 1: cells, Lane 2: Cell free supernatant], (B) Temperature optimization of purified xylanase within 20-100 °C displaying optimum 50 °C, (C) pH optimization between 5.8-7.8 displaying optimum pH 7.0.

**Table 3.** Effect of different media on production of extracellular partially purified xylanase from *Bacillus velezensis* AG20

| Medium | DCW (g/l)<br>*Bacillus*<br>*velezensis* AG20 | Protein (g/l) | Fold<br>Purity | Yield (%) |
| --- | --- | --- | --- | --- |
| TB | 12 | 400 | 5.3 | 21% |
| **AIM** | 8 | 302 | NA | NA |
| **LB** | 3 | 100 | NA | NA |
| **5XLB** | 1 | 75 | NA | NA |

*NA = not measured*

The enzyme activity of partially purified xylanase from *Balillus velezensis* AG20 with 1% (w/v) brichwood xylan statistically optimized on three major factors displayed in table 4. The influence of three factors temperature, pH and buffer ionic concentration were analysed in 27 runs by Taguchi experimental design (Table 5). The response data and S/N ratio of enzyme activity at varying levels are shown in the table 5. Taguchi optimized values for the various optimization parameters are represented in figure 5A, with the prime objective “larger the better” S/N ratio. In the experimental run, the lowest S/N ratio has observed −26.4444 for the combination of 50°C temperature, pH 7 and buffer concentration 50 mM. While, in contrast the combinations 50°C temperature, pH 8 and buffer concentration 70 mM displayed maximum – 12.0412 S/N ratio (Table 5). The best process parameters for highest enzyme activity obtained were the combination of Level 1 parameters 50°C temperature, pH 7 and buffer concentration 50 mM. Main effects of S/N ration on enzyme activity displayed in figure 5A, where buffer ionic concentration showed to have highest effect on enzyme activity as compared to temperature and pH (Figure 5). Linear regression analysis was used to the study of change in enzyme activity (U/ml) at each factor level calculated using following equation:

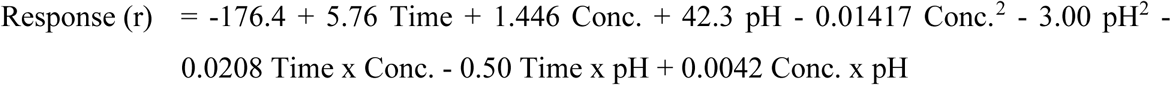

**Figure 5.**
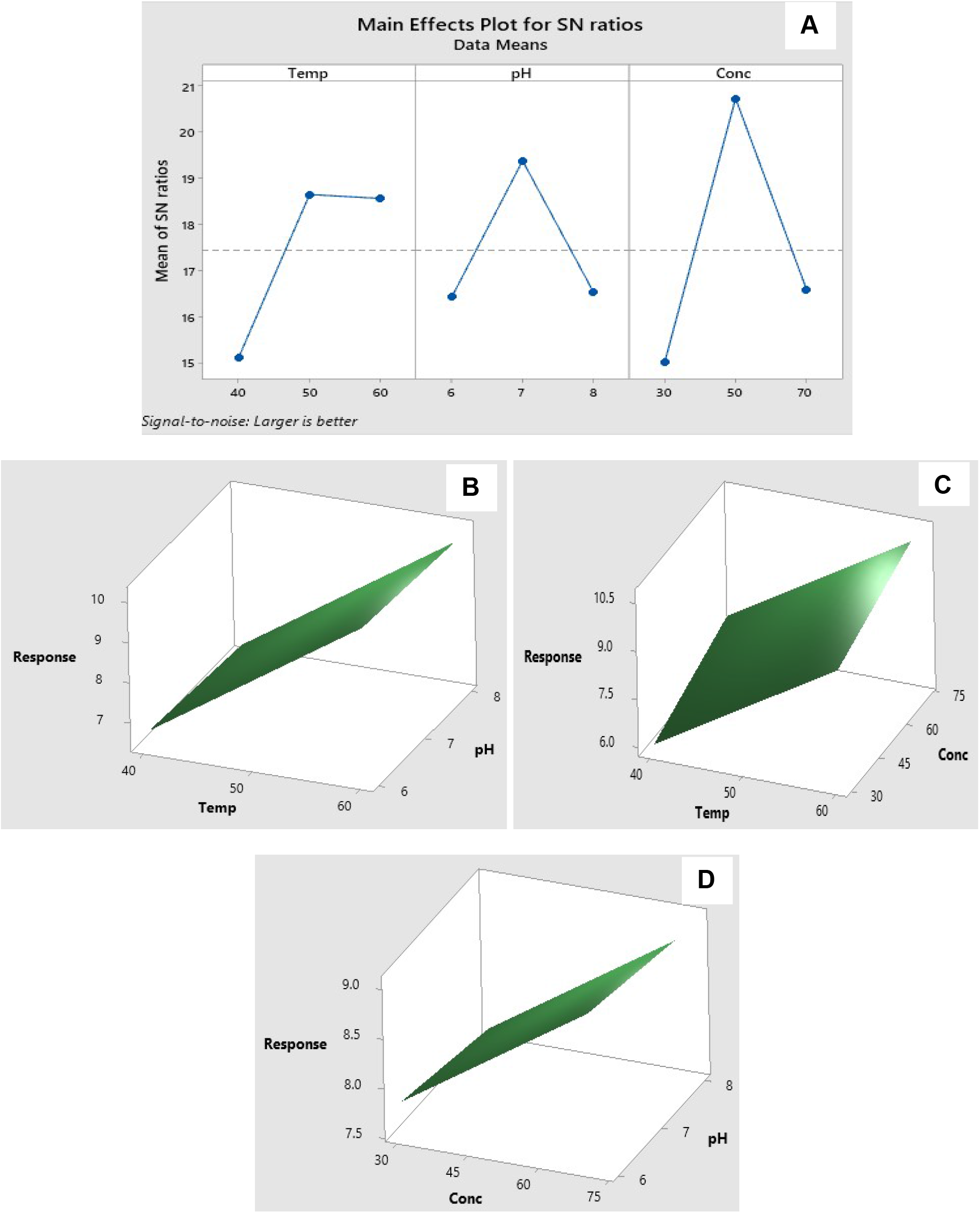
(A) Main effects plots of S/N ratios displaying the effect of each factor under study. Response surface plots for the interaction of (B) pH and temperature, (C) temp and buffer concentration, (D) pH and buffer concentration.

**Table 4.** Optimization parameters and their levels

| **Factors** | **Code** | **Level 1** | **Level 3** | **Level 3** |
| --- | --- | --- | --- | --- |
| **Temperature (°C)** | Temp. | 40 | 50 | 60 |
| **pH** | pH | 6 | 7 | 8 |
| **Buffer ionic concentration (mM)** | Conc. | 30 | 50 | 70 |

**Table 5.** Full factorial design with orthogonal array of Taguchi L-27 (3^3^)

| **Temperature (°C)** | **pH** | **Buffer Conc. (mM)** | **S/N Ratio** |
| --- | --- | --- | --- |
| 40 | 6 | 30 | -13.9794 |
| 40 | 6 | 50 | -13.9794 |
| 40 | 6 | 70 | -13.9794 |
| 40 | 7 | 30 | -13.9794 |
| 40 | 7 | 50 | -15.563 |
| 40 | 7 | 70 | -18.0618 |
| 40 | 8 | 30 | -13.9794 |
| 40 | 8 | 50 | -15.563 |
| 40 | 8 | 70 | -16.902 |
| 50 | 6 | 30 | -13.9794 |
| 50 | 6 | 50 | -25.1055 |
| 50 | 6 | 70 | -15.563 |
| 50 | 7 | 30 | -19.0849 |
| 50 | 7 | 50 | -26.4444 |
| 50 | 7 | 70 | -20.00 |
| 50 | 8 | 30 | -13.9794 |
| 50 | 8 | 50 | -21.5836 |
| 50 | 8 | 70 | -12.0412 |
| 60 | 6 | 30 | -13.9794 |
| 60 | 6 | 50 | -21.5836 |
| 60 | 6 | 70 | -15.563 |
| 60 | 7 | 30 | -18.0618 |
| 60 | 7 | 50 | -25.1055 |
| 60 | 7 | 70 | -13.9794 |
| 60 | 8 | 30 | -13.9794 |
| 60 | 8 | 50 | -18.0618 |
| 60 | 8 | 70 | -16.902 |

Analysis of variance (ANOVA) demonstrated the significance of the model. With R^2^ 90%, R^2^ adjusted 89.64%, P-value 0.0001, high F-value 218.25 and multiple correlation R^2^ 90% indicated the quality of a good model generated using Taguchi’s experimental design at 95% confidence level (significance p<0.05). Moreover, ANOVA has also performed in order to obtain the contribution of each factor influencing enzyme activity. Based on P-value (P<0.05) the contribution of each factor such as temperature (P = 0.0137), pH (P = 0.0232) and buffer concentration (P = 0.004) found significant in influencing the xylanase activity. Interestingly, it was observed that buffer ionic concentration (P = 0.004) played most significant role in maintaining and enhancing the enzyme activity. Interaction profile to the pH and temperature on the enzyme activity displayed high response at pH 7 and temperature 50°C in the surface plot (Figure 5B). Similarly, pH and buffer concentration and temperature and concentration displayed highest response in pH 7 and temperature 50°C and buffer concentration 50 mM (Figure 5C-D). The best possible combination based on P-value suggested at 50°C temperature, pH 7 and phosphate buffer ionic concentration 50 mM highest predicted enzyme activity (Response, (r)) was achieved 20.67 ± 4.0 U/ml. While validating the outcome of Taguchi’s experimental model and the observed enzyme activity 21.0 ± 4.0 found significant for the highest catalsysis of xylan backbone of birchwood xylan.

### ENZYME KINETICS

Kinetic parameters such as K_m_ and V_max_ of the partially purified xylanase were obtained from Michaelis Menten plot (MM plot) and Line Weaver Burk Plot (LB), respectively, as displayed in Table 4, Figure 6A-B. In MM and LB plots the enzyme displayed highest enzyme activity at 1 % (w/v) birch wood xylan was V_max_ = 21.0 ± 3.0 U/ml, K_m_ = 1.25 mg/ml, K_cat_ = 1.75 /s (Table 4, Figure 6A-B). Whereas, the xylanase displayed kinetic values with beech wood xylan V_max_ = 19 ± 2.0 U/ml, K_m_ = 1.1 mg/ml, K_cat_ = 1.58 /s, with glucuronoxylan V_max_ = 11.0 ± 1.0 U/ml, K_m_ = 0.8 mg/ml, K_cat_ = 0.92 /s and with arabinoxylan V_max_ = 4.12 ± 0.4 U/ml, K_m_ = 0.6 mg/ml, K_cat_ = 0.35/s (Table 4).

**Figure 6.**
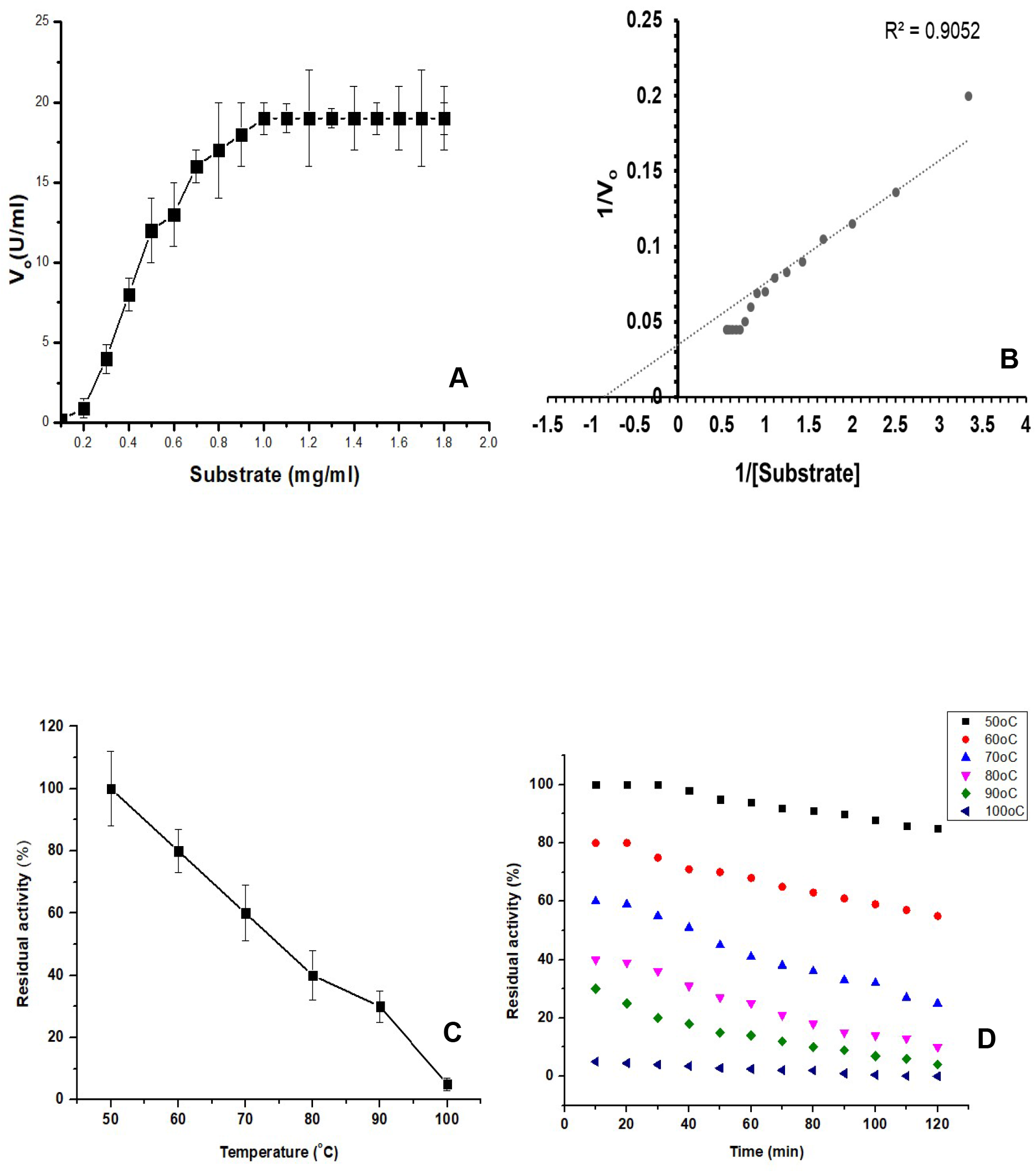
Measurement of enzyme kinetics parameters Vmax, Km and Kcat from (A) hyperbolic Michaelis Menten plot (B) Line weaver Burk plot. Error bars represent mean ± SD. (C) Thermal stability of endoxylanase in elevated temperature from 50 to 100 °C where incubation time is constant 30 min. (D) Time dependent enzyme stability where both the time of incubation (15 – 120 min) and temperature (50–100 °C)

### THERMAL STABILITY STUDY OF THE ENZYME

Thermal stability of the partially purified endoxylanase was examined between 50-100°C. It was appraised from the study that the enzyme was stable between the range 50-60°C and residual activity maintained 80% (Figure 6C). However, at 70, 80, 90 and 100°C residual activity maintained 60%, 40%, 30%, 5%, respectively. Therefore, the sharp fall in residual activity observed from 60-100°C. At 100°C the residual activity was found to be just 5% (Figure 6C). The enzyme was highly stable at 50°C retaining its 100% activity. On the other hand, time dependent thermosability study envisaged the attributes of enzyme at different temperature. It was observed that the endo-xylanase retained it maximum activity (>90%) at 50 °C after 120 min (Figure 6D). However, at 60 °C the retention of residual activity seemed to be less and could be able to retain 55% activity after 120 min. At 70 °C, 80 °C and 90 °C the residual activity fell to 25%, 10% and 4%, respectively (Figure 6D). The enzyme was unable to withstand very high temperature for prolonged period and diminished its entire activity at 100°C after 120 min. Therefore, it can be suggested that the enzyme was moderately thermostable endo-xylanase.

### EFFECT OF METAL IONS AND DETERGENT ON ENZYME ACTIVITY

Enzyme activities are effected by the action metal cation, chelating agents and detergents. Here, the enzyme activity of the endo-xylanase was remarkably increased to 1.5 fold in presence of 5 mM Ca^2+^ ion, 10 mM Mg^2+^ ion. In contrast, Co^2+^, Mn^2+^, Ni^2+^, Zn^2+^, Cu^2+^, Al^3+^ at their very low concentrations drastically reduced the activity of the enzyme (Table 5). With metal cations at 10 mM concentrations of Co^2+^, Mn^2+^, Ni^2+^, Zn^2+^, Cu^2+^, Al^3+^ and Fe^3+^ residual enzyme activity retained was 43%, 65%, 40%, 58%, 30, 39% and 61%, respectively. Whereas, with metal chelating agent EDTA 45% activity was retained. In addition with detergent SDS 10% just residual activity was observed. Under the action of denaturing agents Urea at 1000 mM and GnHCl 10 mM only 5% residual activities were retained (Table 5). The marked decrease in enzyme activity under low concentrations of denaturing agent was due to the denaturation of the enzyme.

### PRODUCTION OF XYLO-OLIGOSACCHARIDE FROM SUGAR CANE BAGGASSE

Sugar cane bagasse is an agro-industrial waste rich in hemicellulose and cellulose content. The efficient pre-treated sugar cane bagasse in FESEM electron micrograph displayed highly porous structure as compared to untreated biomass (Figure 7A-B). Highly porous structure of the biomass gives better space for enzymes to act upon and catalysis leading to efficient biotransformation. Time dependent hydrolysis of pre-treated sugar cane bagasse by partially purified endo-xylanase in TLC displayed the abundance of various degree of polymerized (dp) xylo-oligosaccharides (XOS) (Figure 7C). After 15 min of hydrolysis the initial products obtained were xylose (dp1) and xylobiose (dp2). Then at an interval of 30 min and 1 h of hydrolysis prominent xylose (dp1), xylobiose (dp2) and xylotriose (dp3) spots were observed (Figure 6C). At 6 h interval of hydrolysis higher amount of xylotetrose (dp4) and xylotriose (dp3) were observerd as compared to dp2 and dp1. Similar pattern of hydrolysis were also observed at 12, 18 and 24 h of hydrolysis (Figure 7C). This pattern of random hydrolysis into various degrees of polymerization suggested that the purified enzyme from *Bacillus velezensis* AG20 was an endo-xylanase. The XOS thus produced from the hydrolysis products separated in the pure form from the preparative TLC followed by mass spectroscopy. Mass spectra in positive ionization mode (ESI+) displayed a sharp peak [M+Na]^+^ M/Z = 305.2 for pure xylobiose, [M+Na]^+^ M/Z = 436.5 for xylotriose and [M+Na]^+^ M/Z = 567.0 xylotetrose (Figure 7D-F).

**Figure 7.**
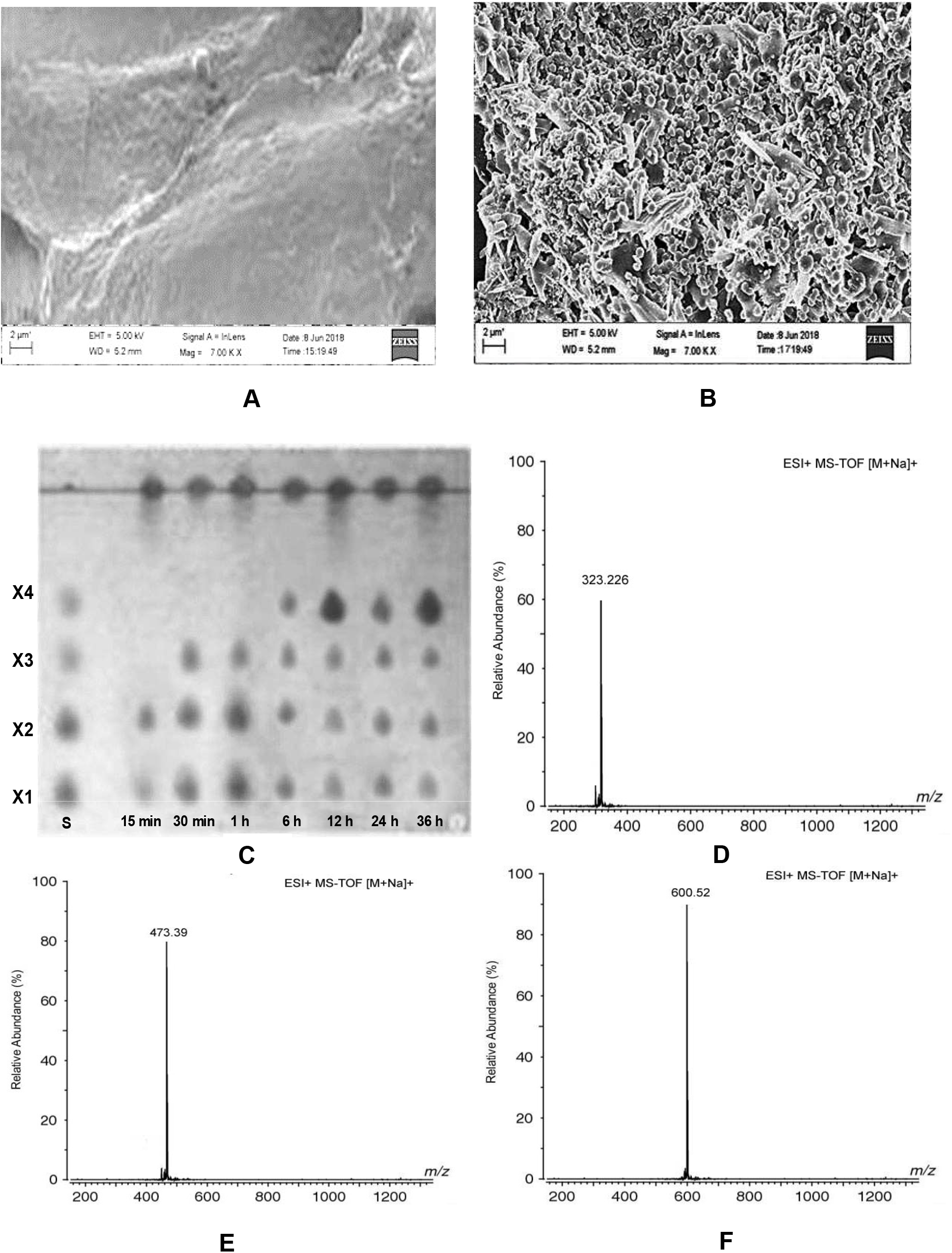
FESEM analysis of sugar cane bagasse (A) before, (B) after pretreatment, (C) TLC of time dependent hydrolyzing (15 min to 36 h) of sugar cane bagasse using partially purified xylanase from *Bacillus velezensis* AG20 displaying the release of XOS of varying degree. Mass spectra peaks in positive ionization mode (ESI+) displayed purified (D) xylobiose, (E) xylotriose and (F) xylotetrose.

### APPLICATION OF MIXED OLIGOSACCHARIDES AS PREBIOTICS AND ANTI-PROLIFIC AGENT

Effect of XOS and inulin as prebiotics in the growth promotion of probiotic bacteria are listed in table 6. Presence of XOS in the culture medium enumerate the augmentation of the growth of probiotics strains *Bifidobacterium infantis* NRRL B-41661, *Bifidobacterium longum* NCC 2705 and *Lactobacillus acidophilus* NRRL B-4495 at 37°C at varying time intervals (Table 6). Mixed XOS promoted the growth of *Bifidobacterium infantis* NRRL B-41661, *Bifidobacterium longum* NCC 2705 and *Lactobacillus acidophilus* NRRL B-4495 from 1×10^7^ cells/ml, 3×10^8^ cells/ml and 1×10^8^ cells/ml to 8×10^7^ cells/ml, 24×10^8^ cells/ml and 9×10^8^ cells/ml, respectively, after 36 h. Whereas, inulin promoted the enumeration upto 4×10^7^ cells/ml, 16×10^8^ cells/ml and 6×10^8^ cells/ml, respectively, after 36 h. A significant ~ 1.5 fold higher growth promotion were observed of the probiotic strains in presence of XOS as compared to commercial prebiotic inulin. By contrast, both XOS and inulin did not support the significant growth promotion of non-probiotic strains *E. coli* DH5α and *E. aerogenes* MTCC 3030 as depicted in table 6.

**Table 7.** Enzyme kinetic parameters of purified xylanase on commercial substrates

| Substrate | V<sub>max</sub> (μmole/min/ml) | K<sub>m</sub> (mg/ml) | K<sub>cat</sub> (/s) |
| --- | --- | --- | --- |
| Birch wood xylan | 21.0 ± 3.0 | 1.25 | 1.75 |
| Beech wood xylan | 19.0 ± 2.0 | 1.10 | 1.58 |
| Glucuronoxylan | 11.0 ± 1.0 | 0.80 | 0.92 |
| Arabinoxylan | 4.12 ± 0.4 U/ml | 0.60 | 0.35 |

**Table 8.** Effect of metal cations, chelating agents, denaturing agents and detergent on the activity of partially purified xylanase from *Bacillus velezensis* AG20

| Reagents/ions | Concentration (mM) | Residual activity (%) |
| --- | --- | --- |
| Ca<sup>2+</sup> | 10 | 150 |
| Mg<sup>2+</sup> | 10 | 150 |
| Co<sup>2+</sup> | 10 | 43 |
| Mn<sup>2+</sup> | 10 | 65 |
| Ni<sup>2+</sup> | 10 | 40 |
| Zn<sup>2+</sup> | 10 | 58 |
| Cu<sup>2+</sup> | 10 | 30 |
| Al<sup>3+</sup> | 10 | 39 |
| Fe<sup>3+</sup> | 10 | 61 |
| EDTA | 10 | 45 |
| SDS | 10 | 10 |
| Urea | 1000 | 5 |
| GnHCl | 10 | 5 |

**Table 9.** Effect of xylo-oligosaccharides (XOS) and commercial inulin on the growth promotion of probiotic and non-probiotic bacteria

| Probiotic and non-probiotic bacteria | Xylo-oligosaccharides (XOS)<br>( x10<sup>7</sup> cells/ml) |  |  |  | Inulin<br>( x10<sup>7</sup> cells/ml) |  |  |  |
| --- | --- | --- | --- | --- | --- | --- | --- | --- |
|  | 0 h | 6 h | 12 h | 36 h | 0 h | 6 h | 12 h | 36 h |
| *Bifidobacterium infantis* NRRL B-41661 | 1 | 4 | 6 | 8 | 1 | 1.5 | 3.1 | 4 |
| *Bifidobacterium longum* NCC 2705 | 30 | 90 | 180 | 240 | 30 | 55 | 134 | 160 |
| *Lactobacillus acidophilus* NRRL B-4495 | 10 | 45 | 70 | 90 | 10 | 30 | 45 | 60 |
| *E. coli* DH5α | 0.2 | 0.4 | 0.8 | 1.1 | 0.2 | 0.4 | 0.7 | 1.2 |
| *Enterobacter aerogenes* MTCC 3030 | 20 | 23 | 27 | 29 | 20 | 22 | 25 | 28 |

In presence of simulated gastric juice (pH 1-4) the mixed XOS displayed significant stability (Figure 8A). XOS displayed 2.5%, 2%, 1.1% and 0.6% of hydrolysis at pH 1, 2, 3 and 4, respectively, after 6 h. However, much higher degree of hydrolysis were observed for inulin 30.1%, 23.6%, 12.8% and 5.6%, respectively, after 6 h of incubation period. Effect of artificial human intestinal fluid displayed significantly low hydrolysis pattern of XOS as compared to inulin (Figure 8B). Percentage of hydrolysis observed for mixed XOS was 1.2%, whereas, commercial inulin degraded in higher degree 6.1% after 6 h of incubation (Figure 8C). This test suggested high stability of XOS in the intestine. Hydrolysis of mixed XOS from pre-treated sugarcane bagasse by α-amylase (pH 5–8) displayed much lower degree of hydrolysis at high pH (Figure 8D). Degree of hydrolysis of XOS were found to be 0.6%, 0.8%, 1.1% and 1.5%, respectively, in pH 5, 6, 7 and 8 after 6 h of incubation. In contrast, inulin displayed higher degree of hydrolysis 9%, 14%, 15.8%, 20%, respectively (Figure 8E). Therefore, the effect of gastric juice, intestinal fluid and enzyme will not hamper much of the integrity of purified XOS from pre-treated sugarcane produced by endo-xylanase activity during their systemic absorption and utilization as prebiotics.

**Figure 8.**
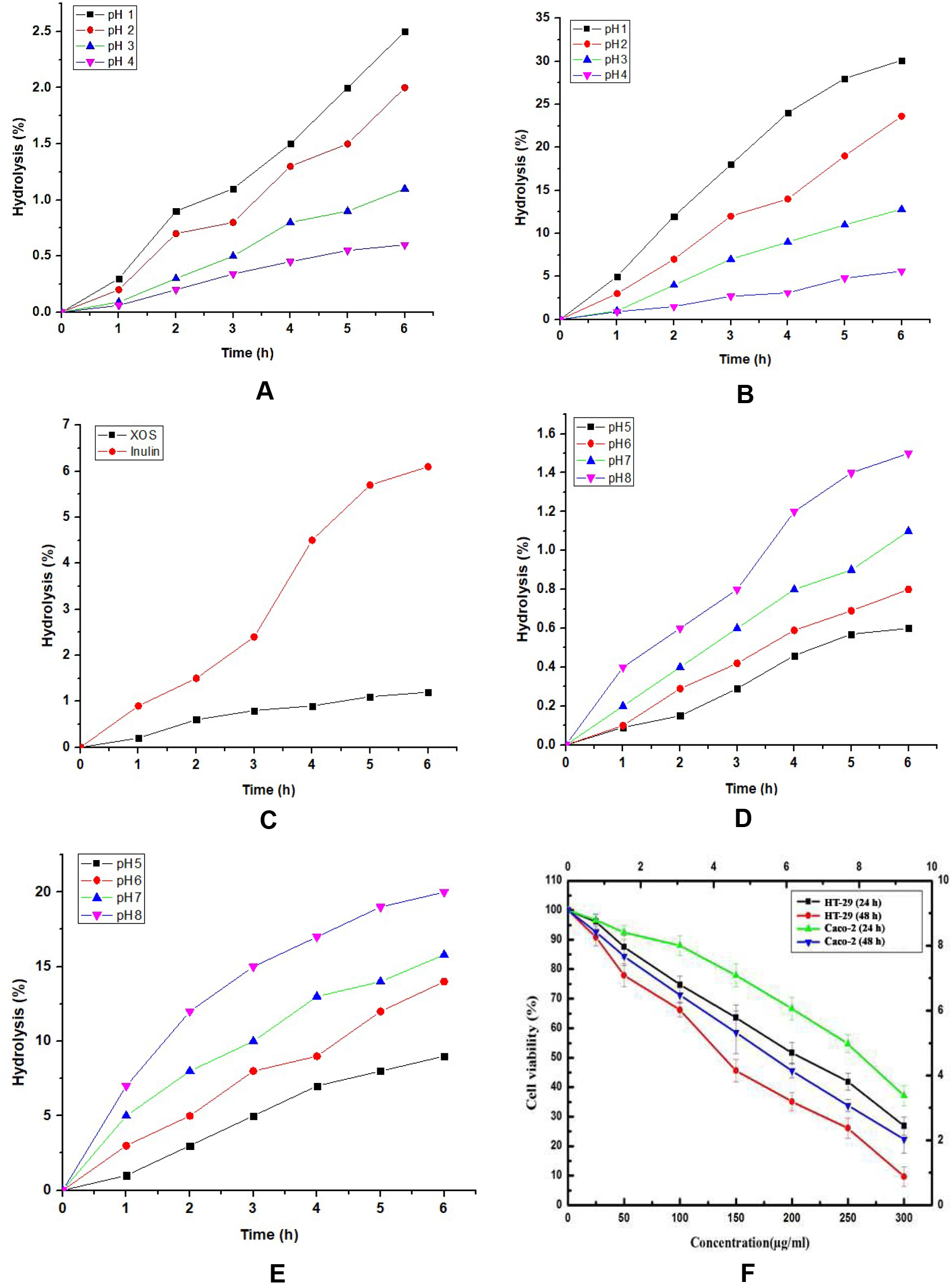
Effect of simulated gastric juice on hydrolysis of (A) mixed xylo-oligosaccharides (XOS) and (B) inulin at pH 1 (purple inverted triangle), 2 (blue up-pointing triangle), 3 (red circle), and 4 (black cubes) at 37 °C for 6 h. (C) Effect of artificial intestinal fluid on hydrolysis of (black cubes) mixed XOS and (red circle) inulin. (D) Effect of α-amylase on the hydrolysis of (D) XOS and (E) inulin at pH 1 (black cubes), pH 2 (red circle), pH 3 (blue up pointed triangle) and pH 4 (purple inverted triangle). (F) mixed XOS (1–300 μg/ mL) on cell viability of human colon carcinoma HT29 and Caco-2 cell lines by MTT assay. Data are expressed as mean ± SD of three independent experiments and each analysis was carried out in triplicate.

In vitro anti-proliferation study of the purified mixed XOS depicted interesting results. In the MTT assay, concentration dependent effect of mixed XOS displayed the significant inhibition of the cell growth of HT-29 and Caco-2 cells (Figure 8F). At low concentration of XOS (300 μg/ml) inhibited 70% and 90% of the cell proliferation of HT-29 cells after 24 h and 48 h, respectively. Nevertheless, the viable cells in case of Caco-2 cell line were found to be 40% and 25%, respectively, after 24 h and 48 h of treatment at 300 μg/ml concentration of mixed XOS (Figure 7F). These results suggested that XOS has high anti-proliferation activity against colon cancer cells.

## DISCUSSIONS

Cave is the extreme environment of nature and a reservoir for novel extremophilic microoganisms that lead their sustainability by chemical conversion. Cultural, biochemical and molecular identification facilitate the determination of bacterial species and taxonomic affirmations form the unknown sample (11, 41). In our present study, we have determined the bacterial species isolated from the soil sample of Krem Phyllut cave of Meghalaya. Out of nine isolated colonies, three colonies were depicted gram positive and efficient carbohydrate fermenter. These outcomes corroborate with the salient features of the genera *Bacillus*. In vitro antibiotic susceptibility test found resistance to the broad spectrum antibiotics that corroborates with the findings about the ancient bacterium *Paenibacillus* isolated from the deepest Lechuguilla Cave, New Mexico resistant to the most powerful antibiotics of modern medicine (42). They have also suggested that the antibiotic resistance characteristics led this organism into a superbug family. 16S rDNA sequencing based molecular, phylogenetic analysis and RNA secondary structure folding energy let to identify the strain producing maximum extracellular endo-xylanase was predominantly a *Bacillus velezensis* AG20. This is the first ever report of *Bacillus velezensis* from the caves of Meghalaya, India and rest of the world ever. Earlier, this novel group of bacteria was reported from plant root isolate and nutrient poor desert soil secretes beneficial metabolites for plant growth promotion and protection as probiotics (43, 44). Positive correlation of high GC content and basic melting temperature were owing to thermotolerent property of *Bacillus velezensis* AG20. Similarly, high and optimal GC content of prokaryotes led to the high thermal adaptability was also reported elsewhere (45). The evidence of biofilm formation in electron micrograph by of the novel *Bacillus velezensis* AG20 was also a feature of cave microbes similarly reported earlier by Banerjee and coworker (2, 7). Thermostable and thermophilic xylanases have much industrial significance. There is a need for efficient method for the production of high level of xylanase from *Bacillus velezensis* AG20. Therefore, several medium were tested for higher production extracellular xylanase. It was found that medium composition played a significant role where enriched medium terrific broth (TB) supported high production of cell biomass as well as primary metabolite xylanase. Similar observations were portrayed by Lu et al., (2017), for higher production of extracellular recombinant protein from *Bacillus cereus*. TB promotes higher cell growth and due to the phosphate buffer action in the medium composition facilitates controlled higher extracellular enzyme production. The extracellular xylanase thus purified exhibited significant enzyme activities at optimum pH and temperature 7 and 50°C, respectively. Similar observations were also reported where *Bacillus licheniformis* DM5 displayed high enzyme activities at similar pH and temperature (11). Moreover, retention of 20% enzyme activity in alkaline condition may facilitate the utilization of this enzyme in industry. The enzyme activity under variable pH, temperature and buffer ionic concentrations found to be statistically significant. The choice of the substrate for enzymatic catalysis, the purified extracellular xylanase showed unique characteristics. Linear chain of xylan polymer was found to be suitable for catalysis *viz*. birchwood xylan, beechwood xylan linked via β-(1→4)-glycosidic linkages between the monomers. Confronting the hydrolysis of appendaged polysaccharides such as glucuronoxylan and arabinoxylan, which are moderately and highly branched, respectively. Although, the purified xylanase exhibited high activity against linear xylan polymers but also exhibit low levels of appendage dependent catalysis of branched polymers. Similar observations were depicted in a study where *Bacillus licheniformis* DM5 showed maximum likeliness to our findings (11). Nevertheless, contrasting activity against pNP-xylopyranoside ruled out the possibility of having exo-xylanase or any xylosidase activity. The action pattern of the xylanase enzyme belongs to the glycoside hydrolase family 30 (XynGH30) represent novel properties of hydrolysis. Enzyme activity can be augmented by the presence of cofactors Ca^2+^ and Mg^2+^ ions. Similar report envisaged in an earlier study where thermostable enzyme activity enhanced 1.5 fold by Ca^2+^ and Mg^2+^ ions (47). Moreover, the enzyme depicted moderately thermostable which would have high demand in the industry. Linear and appendage dependent studies involved in xylo-oligosaccrides (XOS) production from xylan rich substrates also corroborated with the earlier findings described elsewhere (11, 48, 49, 50, 51). XOS produced from agro-industrial waste sugar cane bagasse improved the growth of probiotic strains as potential prebiotics. Nevertheless, XOS thus produced from the agro waste displayed higher order of stability under the action of gastric juice, intestinal fluid and α-amylase upto 6 hr. Apparently, 90% of the XOS will remain intact under the systemic factors influence for better performance as prebiotics. This result corroborated with the earlier report (34). The discrimination in growth promotion by mixed xylo-oligosaccharides of probiotic *Lactobacillus* and *Bifidobacterium* was due to the generation of short chain fatty acids and lactate from anaerobic fermentation in the presence of prebiotics that involved in supply energy for colonic epithelium and suppression of pathogenic intestinal bacteria by lowering of pH (35). These findings corroborated the earlier reports (52, 53). which explained the growth of *Lactobacillus* sp. and *Bifidobacterium* sp. in low pH. *In vitro* anti-proliferation assay of mixed XOS against colon cancer cell lines HT-29 and Caco-2 were the novel findings in the present study. At low concentrations (300 μg/ml) of mixed XOS suppressed the proliferation of HT-29 and Caco-2 cells 90% and 75%, respectively, after 48 h. Such a finding can initiate the advancement of application of XOS of this kind in anticancer therapy and may pave the applications of XOS as nutraceutical and theragonostic implications as well. Earlier studies suggested that the fructooligosaccharides and manno-oligosaccharides from apple and coconut were effective against human colon cancer cell line due to cell cycle arrest in S phase leading to apoptosis (34, 54). Similar mechanisms might have enlighten in the present study. Therefore, thermostable xylanase from the novel strain *Bacillus velezensis* AG20 isolated from the cave of Meghalaya, India and its applications in XOS production may surpass the demand for future enzyme, nutraceutical, prebiotic for industry and sustainable biofuel.

## Acknowledgement

Authors would like to acknowledge the Central Instrumentation Facility, Gauhati University for the FESEM studies. This research received no specific grant from any funding agency in the public, commercial, or not for profit sectors.

## Author’s contribution

Experimental procedures were designed and carried out by DB, AG, SM and SB. Manuscript preparation was done by AG and DB.

## Compliance with ethical standards

This study does not contain any studies with animals performed by any of the authors

## REFERENCES

1. Daly BDK. 2009. Meghalaya’s underground treasures, in Glimpses from the North-East: National Knowledge Commission, pp 49–54.

2. Banerjee S, Rai S, Sarma B, Joshi SR. 2012. Bacterial biofilm in water bodies of Cherrapunjee: the rainiest place on planet earth. Adv Microbiol. 2:465–475.

3. Sarbu MS, Kane CT, Kinkle KB. 1996. A Chemoautotrophically Based Cave Ecosystem. Science. 272(5270):1953–1955.

4. Northup ED, Lavoie HK. 2010. Geomicrobiology of caves: a review. Geomicrobiol J. 18(3): 199–222.

5. Baskar S, Baskar R., Lee N, Theophilus PK. 2009. Speleothems from Mawsmai and Krem Phyllut caves, Meghalaya, India: some evidences on biogenic activities. Environ. Geol, 57(5): 1169–1186

6. Mudgil D, Sushmitha B, Ramanathan BD, Paulcand Y, Shouche S. 2018. Biomineralization Potential of *Bacillus subtilis, Rummelii bacillus* Stabekisii and *Staphylococcus epidermidis* Strains in vitro isolated from speleothems, Khasi Hill Caves, Meghalaya, India. Geomicrobiol J. DOI: 10.1080/01490451.2018.1450461

7. Banerjee S, Joshi SR. 2016. Culturable bacteria associated with the caves of meghalaya in india contribute to speleogenesis. J Cave Karst Stud. 78(3): 144–157

8. Riding R. 2000. Microbial carbonates: the geological record of calcified bacterial-algal mats and biofilms: Sedimentology. 47:179–214

9. Banka AL, Guralp SA, Gulari E. 2014. Secretory expression and characterization of two hemicellulases, xylanase, and β-xylosidase, isolated from *Bacillus subtilis* M015. Appl Biochem Biotechnol. 174(8):2702–2710

10. Thomas L, Ushasree MV, Pandey A. 2014. An alkali-thermostable xylanase from Bacillus pumilus functionally expressed in *Kluyveromyces lactis* and evaluation of its deinking efficiency. Bioresour Technol. 165:309–313

11. Ghosh A, Sutradhar S, Baishya D. 2019. Delineating thermophilic xylanase from *Bacillus licheniformis* DM5 towards its potential application in xylooligosaccharides production. World J Microbiol Biotechnol, 35(34):https://doi.org/10.1007/s11274-019-2605-1

12. Bernier JR, Driguez H, Desrochers M. 1983. Molecular cloning of a *Bacillus subtilis* xylanase gene in *Escherichia coli*. Gene. 26(1):59–65.

13. Elegir G, Szakács G, Jeffries TW. 1994. Purification, characterization, and substrate specificities of multiple xylanases from *Streptomyces* sp. Strain B-12-2. Appl Environ Microbiol. 60(7):2609–2615.

14. Collins T, Hoyoux A, Dutron VA, Georis J, Genot B, Dauvrin T, Amaut F, Gerday C, Feller G. 2006. Use of glycoside hydrolase family 8 xylanases in baking. J. Cereal Sci. 43(1):79–84.

15. Singh YD, Satapathy K.B. 2018. Conversion of lignocellulosic biomass to bioethanol: an overview with a focus on pretreatment. Int J Engg Technol. 15:17–43.

16. Gupta S, Kuhad RC, Bhushan B., Hoondal GS. 2001. Improved xylanase production from a haloalkalophilic *Staphylococcus* sp. SG-13 using inexpensive agricultural residues. World J Microbiol Biotechnol. 17(5):https://doi.org/10.1023/A:1016691205518

17. Dwivedi P, Vivekanand V, Ganguly R, Singh RP. 2009. *Parthenium* sp. as a plant biomass for the production of alkali tolerant xylanase from mutant *Penicillium oxalicum* SAU_E_-3.510 in submerged fermentation. Biomass Bioener. 581–588.

18. Chugh P, Soni R. Soni SK. 2016. Deoiled rice bran: a substrate for co-production of a consortium of hydrolytic enzymes by *Aspergillus niger* P-19. Waste Biomass Valori. 7(513): https://doi.org/10.1007/s12649-015-9477-x

19. Maciel GM, Vandenberghe LS, Windson C, Haminiuk I, Fendrich RC, Bianca BD, Brandalize TQ, Pandey A, Soccol CR. 2008. Xylanase production by *Aspergillus niger* LPB 326 in solid-state fermentation using statistical experimental designs. Food Technol Biotechnol. 46(2):183–189.

20. Javier A. Linares P, Anna A, Eva NK. 2018. Structural considerations on the use of endo-xylanases for the production of prebiotic xylooligosaccharides from biomass. Curr Protein Pept Sc. 19(1):48–67.

21. Wang J, Sun B, Cao Y, Tian Y, Wang C. 2009. Enzymatic preparation of wheat bran xylooligosaccharides and their stability during pasteurization and autoclave sterilization at low pH. Carbohydr Polym. 77:816–821.

22. Verma D, Anand A, Satyanarayana T. 2013. Thermostable and alkalistable endoxylanase of the extremely thermophilic bacterium *Geobacillus thermodenitrificans* TSAA1: Cloning, expression, characteristics and its applicability in generating xylooligosaccharides and fermentable sugars. Appl Biochem Biotechnol. 13(170):119–130.

23. Mandelli F, Brenelli L.B, Almeida R.F, Goldbeck R, Wolf LD, Hoffmam ZB, Ruller R, Rocha GJ, Mercadante AZ, Squina FM. 2014. Simultaneous production of xylooligosaccharides and antioxidant compounds from sugarcane bagasse *via* enzymatic hydrolysis. Ind Crops Prod. 14(52):770–775.

24. Faryar R., Linares-Pastén JA, Immerzeel P, Mamo G, Andersson M, Stålbrand H, Mattiasson B, Nordberg KE. 2015. production of prebiotic xylooligosaccharides from alkaline extracted wheat straw using the k80r-variant of a thermostable alkali-tolerant xylanase. Food Bioprod Process. 93:1–10.

25. Gibson GR, Roberfroid MB. 1995. Dietary modulation of the human colonic microbiota: introducing the concept of prebiotics. J Nutr. 125(6):1401–1412.

26. Christakopoulos P, Katapodis P, Kalogeris E, Kekos D, Macris BJ, Stamatis H, Skaltsa H. 2003. Antimicrobial activity of acidic xylo-oligosaccharides produced by family 10 and 11 endoxylanases. J Biol Macromol. 31(4-5):171–175.

27. Barreteau H, Delattre C, Michaud P. 2006. Production of oligosaccharides as promising new food additive generation. oligosaccharides as food additives. Food Technol Biotechnol. 44(3):323–333.

28. Cappuccino JG, Sherman N. 1996. Microbiology - A Laboratory Manual, The Benjamin/Cummings Publishing Co., Inc., Menlo Park, California.

29. Bauer AW, Kirby WMM, Sherris JC, Turck M. 1966. Antibiotic susceptibility testing by a standardized single disk method. Am J Clin Pathol. 36:493–496.

30. Johnsen H, Krause K. 2014. Cellulase activity screening using pure caroxymethyl cellulose: application to soluble cellulolytic samples and to plant tissue prints. Int J Mol Sci 15:830–838

31. Zuker M. 2003. Mfold web server for nucleic acid folding and hybridization prediction. Nucleic Acids Res. 31(13):3406–3415

32. Tripathi NK, Shrivastva A, Biswal KC, Rao PVL. 2009. Optimization of culture medium for production of recombinant dengue protein in *Escherichia coli*. Ind Biotechnol 5:179–183

33. Kim S. 2018. Enhancing Bioethanol Productivity Using Alkali-Pretreated Empty Palm Fruit Bunch Fiber Hydrolysate. BioMed Res Int. https://doi.org/10.1155/2018/5272935

34. Ghosh A, Verma A, Rao JMT, Shukla R, Goyal A. 2015. Recovery and purification of oligosaccharides from copra meal by recombinant endo-β-mannanase and deciphering molecular mechanism involved and its role as potent therapeutic agent. Mol Biotechnol. 57:111–127

35. Hongpattarakere, T. 2013. Improvement of freeze-dried *Lactobacillus plantarum* survival using water extracts and crude fibers from food crops. Food Bioproc Tech. 6:1885–1896.

36. Korakli M, Ganzle MG, Vogel RF. 2002. Metabolism by *Bifidobacteria* and lactic acid bacteria of polysaccharides from wheat and rye, and exopolysaccharides produced by *Lactobacillus sanfranciscensis*. J Appl Microbiol. 92:958–965

37. Nelson N. 1944. A photometric adaptation of the Somogyi method for the determination of glucose. J Biol Chem. 153:375–380.

38. Somogyi M. 1945. A new reagent for the determination of sugars. J Biol Chem 160:61–68.

39. Wichienchot S, Jatupornpipat M, Rastall RA. 2010. Oligosaccharides of pitaya (dragon fruit) flesh and their prebiotic properties. Food Chem. 120:850–857.

40. Mosmann T. 1983. Rapid colorimetric assay for cellular growth and survival: Application to proliferation and cytotoxicity assays. J Immunol Methods. 65:55–63.

41. Harley JP. 2008. Laboratory exercises in microbiology, 7th edn. McGraw-Hill Companies, New York.

42. Bhullar K, Waglechner N, Pawlowski A, Koteva K, Banks ED, Johnston MD, Barton HA, Wright GD. 2012. Antibiotic resistance is prevalent in an isolated cave microbiome. PloS one. 7(4):e34953.

43. Rabbee MF, Ali MS, Choi J, Hwang BS, Jeong SC, Baek KH. 2019. *Bacillus velezensis:* a valuable member of bioactive molecules within plant microbiomes. Molecules. 24(6):1046: doi:10.3390/molecules24061046

44. Reva ON, Dirk ZHS, Liberata AM, Aneth DM, Dillon M, Monique J, Wai YC, Stefanie L, Christian HA, Lylia VA, Maksim AK, Donatha T, Sylvester L, Joachim V, Rainer B, Johan Meijer. 2019. Genetic, epigenetic and phenotypic diversity of four *bacillus velezensis* strains used for plant protection or as probiotics. Front Microbiol. https://doi.org/10.3389/fmicb.2019.02610

45. Basak S, Mukhopadhyay P, Gupta SK, Ghosh TC. 2010. Genomic adaptation of prokaryotic organisms at high temperature. Bioinformation. 4(8):352–356.

46. Lu S, Jiang Q, Yu L, Wu J. 2017. Enhanced extracellular production of recombinant proteins in Escherichia coli by co-expression with *Bacillus cereus* phospholipase C. Microb Cell Fact 16(1):doi:10.1186/s12934-017-0639-3

47. Ghosh A, Luis AS, Brás JLA, Fontes CMGA, Goyal A. 2013. Thermostable recombinant endo-β-(1 → 4)-mannanase from *Clostridium thermocellum:* Biochemical characterization and manno-oligosaccharides production J Agric Food Chem. 61:12333–12344.

48. Biely P, Vrsanska M, Kremnicky L, Tenkanen M, Poutanen K, Hayn M. 1993. Catalytic properties of endo-β-1,4-xylanases of *Trichoderma reesei*. In: Suominen P, Reinikainen T (eds) Trichoderma reesei cellulases and other hydrolases, Fagepaino Oy, Helsinki, pp. 125–135

49. Hurlbert JC, Preston JF. 2001. Functional characterization of a novel xylanase from a corn strain of *Erwinia chrysanthemi*. J Bacteriol 183:2093–2100

50. Saloheimo M, Siika-aho M, Tenkanen M, Penttila ME. 2003. Novel xylanase from *Trichoderma reesei*, method for production thereof, and methods employing this enzyme. United States Patent Application 20030054518

51. Tenkanen M, Burgermeister M, Vrsanska M, Biely P, Saloheimo M, Siika-aho M. 2003. A novel xylanase XYN IV from *Trichoderma reesei* and its action on different xylans. In: Courtin CM, Veraverbeke WS, Delcour JA (eds) Recent advances in enzymes in grain processing. The Katholieke Universiteit Leuven, Leuven, pp 41–46

52. Gibson GR, Ottaway PB, Rastall RA. 2000. Prebiotics: new developments in functional foods. Wood head Publishing Limited, Oxford

53. Pereira DIA, Gibson GR (2002) Cholesterol assimilation by lactic acid bacteria and Bifidobacteria isolated from the human gut. Appl Environ Microbiol 68:4689–4693

54. Li Q, Zhou S, Jing J, Yang T, Duan S, Wang Z, Mei Q, Liu L (2013) Oligosaccharide from apple induces apoptosis and cell cycle arrest in HT29 human colon cancer cells. Int J Biol Macromol 57:245–254

